# Differences in the gut microbiomes of distinct ethnicities within the same geographic area are linked to host metabolic health

**DOI:** 10.1101/2020.10.23.352807

**Authors:** Qi Yan Ang, Diana L. Alba, Vaibhav Upadhyay, Jordan E. Bisanz, Jingwei Cai, Ho Lim Lee, Eliseo Barajas, Grace Wei, Cecilia Noecker, Andrew D. Patterson, Suneil K. Koliwad, Peter J. Turnbaugh

**Affiliations:** Department of Microbiology and Immunology, G.W. Hooper Research Foundation, University of California, San Francisco, CA 94143, USA; Diabetes Center, University of California San Francisco, CA 94143, USA; Division of Endocrinology, Diabetes, and Metabolism, Department of Medicine, University of California San Francisco, CA 94143, USA; Center for Molecular Toxicology and Carcinogenesis, Department of Veterinary & Biomedical Sciences, Pennsylvania State University, PA 16802, USA

**Keywords:** human gut microbiome, ethnicity, multi-omics, metabolic syndrome, obesity, biogeography

## Abstract

**Background:** The human gut microbiota exhibits marked variation around the world, which has been attributed to dietary intake and other environmental factors. However, the degree to which ethnicity-associated differences in gut microbial community structure and function are maintained following immigration or in the context of metabolic disease is poorly understood.

**Results:** We conducted a multi-omic study of 46 lean and obese East Asian and White participants living in the San Francisco Bay Area. 16S rRNA gene sequencing revealed significant differences between ethnic groups in bacterial richness and community structure. White individuals were enriched for the mucin-degrading *Akkermansia muciniphila.* East Asian participants had increased levels of multiple bacterial phyla, fermentative pathways detected by metagenomics, and the short-chain fatty acid end products acetate, propionate, and isobutyrate. Differences in the gut microbiota between the East Asian and White groups could not be explained by reported dietary intake, were more pronounced in lean individuals, and were associated with current geographical location. Microbiome transplantations into germ-free mice confirmed that the differences in the gut microbiota of the East Asian and White individuals we analyzed are independent of diet and that they differentially impact host body weight and adiposity in genetically identical mouse recipients.

**Conclusions:** The reported findings emphasize the utility of studying diverse ethnic groups within a defined geographical location and provide a starting point for dissecting the mechanisms contributing to the complex interactions between the gut microbiome and ethnicity-associated lifestyle, demographic, metabolic, and genetic factors.

## BACKGROUND

Culture-independent surveys have emphasized differences in gut microbial community structure between countries [1–3]; however, the factors that contribute to these differences are poorly understood. Diet is a common hypothesis for geographical variations in the gut microbiota [4, 5], based upon extensive data from intervention studies in humans and in mouse models [6–9]. However, diet is just one of the many factors that distinguishes human populations at the global scale, motivating the desire for a more holistic approach to identify determinants of microbiota composition. Self-identified race/ethnicity (SIRE) provides a useful alternative, as it integrates the broader national or cultural tradition of a given social group and is closely tied to both dietary intake and genetic ancestry. Associations between the gut microbiota and ethnicity have been reported in China [10], the Netherlands [11], Singapore [12], and the United States [13, 14]. In contrast, a recent study of Asian immigrants suggested that once an individual relocates to a new country, the microbiota rapidly assumes the structure of the country of residence [3]. Thus, the degree to which microbiome signatures of ethnicity persist following immigration and their consequences for host pathophysiology remain an open question.

The links between ethnicity and metabolic disease are well-established. For example, East Asian (EA) subjects are more likely to develop health-related metabolic complications at lower body mass index (BMI) compared to their White (W) counterparts [15, 16]. Moreover, Asian Americans have persistent ethnic differences in metabolic phenotypes following immigration [17], including a decoupling of BMI from total body fat percentage [18]. The mechanisms contributing to these ethnic differences in fat accrual and its metabolic impact remain unknown. Human genetic polymorphisms may play a role [19, 20]; however, putative alleles are often shared between members of different ethnic groups [21]. Alternatively, the gut microbiome might offer a possible explanation for differences in metabolic disease rates across ethnic groups, but there has been a relative scarcity of microbiome studies in this area.

These observations led us to hypothesize that ethnicity-associated differences in host metabolic phenotypes may be determined by corresponding differences in the gut microbiome. First, we sought to better understand the extent to which ethnicity is linked to the human gut microbiome in states of health and disease. We conducted a cross-sectional multi-omic analysis of the gut microbiome using paired 16S rRNA gene sequencing (16S-seq), metagenomics, and metabolomics from the Inflammation, Diabetes, Ethnicity, and Obesity (IDEO) cohort at the University of California, San Francisco. IDEO includes rich metabolic, dietary, and socioeconomic metadata [18], a restricted geographical distribution within the San Francisco Bay Area, and a balanced distribution of EA and W individuals that are both lean and obese (**Table S1**). We report marked differences in gut microbial richness, community structure, and metabolic end products between EA and W individuals in the IDEO cohort. We then used microbiome transplantations to assess the stability of ethnicity-associated differences in the gut microbiota in the context of genetically identical mice fed the same diet. We also explored the functional consequences of these differences on host metabolic phenotypes. Our results emphasize the importance of considering ethnicity in microbiome research and further complicate prior links between metabolic disease and the gut microbiome [22–24], which may be markedly different across diverse ethnic groups.

## METHODS

### Human subjects

All study participants were part of the IDEO cohort, which has been previously described [18, 25]. Briefly, IDEO consists of 25-65 year-old men and women of multiple ethnicities and across a wide BMI range (18.5–52 kg/m2) living in the San Francisco Bay Area; exclusion factors include smoking, unstable weight within the last 3 months (>3% weight gain or loss), a diagnosed inflammatory or infectious disease, liver failure, renal dysfunction, cancer, and reported alcohol consumption of >20 grams per day. Using IDEO, we recruited both lean and obese W and EA individuals into this study based on World Health Organization cut-offs: W/EA BMI**≤**24.9 kg/m^2^ (lean); W BMI≥30 kg/m^2^ (obese); and EA BMI≥27.5 kg/m^2^ (obese) [17,26,27].

Each participant consented to take part in the study, which was approved by the University of California San Francisco (UCSF) Committee on Human Research. We utilized demographic, medical, dietary, and lifestyle metadata on each participant that were part of their initial recruitment into IDEO, as previously reported [18,25,28]. Participants with Type 2 Diabetes (T2D) were classified in accordance with American Diabetes Association Standards of Medical Care guidelines [29], defined by having glycated hemoglobin (HbA1c) ≥6.5% or the combination of a prior diagnosis of T2D and the active use of an antidiabetic medication. For stool sample collection, participants took home or were mailed a stool sample collection kit and detailed instructions on how to collect the specimen. All samples were collected at home, stored at room temperature, and brought to the UCSF Clinical Research Center by the participants within 24 hours of defecation. Samples were aliquoted and stored at -80°C.

### Anthropometric and body composition measurements

We leveraged host phenotypic and demographic data from IDEO, which was the focus of two previous studies [18, 25]. For the convenience of the reader, we restate our methods here. Height and weight were measured using a standard stadiometer and scale, and BMI (kg/m^2^) was calculated from two averaged measurements. Waist and hip circumferences (to the nearest 0.5 cm) were measured using a plastic tape meter at the level of the umbilicus and of the greater trochanters, respectively, and waist-to-hip ratio (WHR) was calculated. Blood pressure was measured with a standard mercury sphygmomanometer on the left arm after at least 10 minutes of rest. Mean values were determined from two independent measurements. Blood samples were collected after an overnight fast and analyzed for plasma glucose, insulin, serum total cholesterol, high density lipoprotein (HDL) cholesterol, and triglycerides. Low density lipoprotein (LDL) cholesterol was estimated according to the Friedewald formula [30]. Insulin resistance was estimated by the homeostatic model assessment of insulin resistance (HOMA-IR) index calculated from fasting glucose and insulin values [31]. Two obese subjects on insulin were included in the HOMA-IR analysis (1 EA, 1 W). Body composition of the subjects was estimated by Dual-Energy X-ray Absorptiometry (DEXA) using a Hologic Horizon/A scanner (3-minute whole-body scan, <0.1 G milligray) per manufacturer protocol. A single technologist analyzed all DEXA measurements using Hologic Apex software (13.6.0.4:3) following the International Society for Clinical Densitometry guidelines. Visceral adipose tissue (VAT) was estimated from a 5 cm-wide region across the abdomen just above the iliac crest, coincident with the fourth lumbar vertebrae, to avoid interference from iliac crest bone pixels and matching the region commonly used to analyze VAT mass by CT scan [32–34] . The short version of the International Physical Activity Questionnaire (IPAQ) was used to assess the habitual physical activity levels of the participants. The IPAQ total score is expressed in metabolic equivalent (MET)-minutes/week [35].

### Dietary assessment

IDEO participants completed two dietary questionnaires, as previously described [18, 25], allowing for the assessment of usual total fiber intake and fiber from specific sources, as well as macronutrient, phytochemical, vitamin, and mineral uptake. The first instrument was an Automated Self-Administered 24-hour Dietary Assessment (ASA24) [36, 37], which queries intake over a 24-hour period. The 24-hour recalls and supplement data were manually entered in the ASA24 Dietary Assessment Tool (v. 2016), an electronic data collection and dietary analysis program. ASA24 employs research-based strategies to enhance dietary recall using a respondent-driven approach allowing initial recall to be self-defined. The second instrument was the National Cancer Institute’s Diet History Questionnaire III (DHQIII) [38, 39]. The DHQIII queries one’s usual diet over the past month [39]. Completing the DHQIII was associated with participant survey fatigue and completion rates were accordingly only 42% after 1 phone-based administration of the instrument, although they improved to 79% by the 2nd session and reached 100% within four sessions over a 5-month period. Due to the effort needed to achieve DHQIII completion, we modified our protocol to request completion of the simpler ASA24 at three separate times, at appointments where there were computers and personnel assistance for online completion, in addition to completion of the DHQIII questionnaire. By combining both instruments, we were able to reliably obtain complete dietary information on all participants.

### DNA extraction

Human stool samples were homogenized with bead beating for 5 min (Mini-Beadbeater-96, BioSpec) using beads of mixed size and material (Lysing Matrix E 2mL Tube, MP Biomedicals) in the digestion solution and lysis buffer of a Wizard SV 96 Genomic DNA kit (Promega). The samples were centrifuged for 10 min at 16,000 g and the supernatant was transferred to the binding plate. The DNA was then purified according to the manufacturer’s instructions. Mouse fecal pellets were homogenized with bead beating for 5 min (Mini-Beadbeater-96, BioSpec) using the ZR BashingBead lysis matrix containing 0.1 and 0.5 mm beads (ZR-96 BashingBead Lysis Rack, Zymo Research) and the lysis solution provided in the ZymoBIOMICS 96 MagBead DNA Kit (Zymo Research). The samples were centrifuged for 5 min at 3,000 g and the supernatant was transferred to 1 mL deep-well plates. The DNA was then purified using the ZymoBIOMICS 96 MagBead DNA Kit (Zymo Research) according to the manufacturer’s instructions.

### 16S rRNA gene sequencing and analysis

For human samples, 16S rRNA gene amplification was carried out using GoLay-barcoded 515F/806R primers [40] targeting the V4 region of the 16S rRNA gene according to the methods of the Earth Microbiome Project (earthmicrobiome.org) (**Table S2**). Briefly, 2 µL of DNA was combined with 25 µL of AmpliTaq Gold 360 Master Mix (Fisher Scientific), 5 µL of primers (2 µM each GoLay-barcoded 515/806R), and 18 µL H_2_O. Amplification was as follows: 10 min 95°C, 30x (30s 95°C, 30s 50°C, 30s 72°C), and 7 min 72°C. Amplicons were quantified with PicoGreen (Quant-It dsDNA; Life Technologies) and pooled at equimolar concentrations. Aliquots of the pool were then column (MinElute PCR Purification Kit; Qiagen) and gel purified (QIAquick Gel Extraction Kit; Qiagen). Libraries were then quantified (KAPA Library Quantification Kit; Illumina) and sequenced with a 600 cycle MiSeq Reagent Kit (250×150; Illumina) with ∼15% PhiX spike-in. For mouse samples, 16S rRNA gene amplification was carried out as per reference protocol and primers [41]. In brief, the V4 region of the 16S rRNA gene was amplified with 515F/806R primers containing common adaptor sequences, and then the Illumina flow cell adaptors and dual indices were added in a secondary amplification step (see **Table S3** for index sequences). Amplicons were pooled and normalized using the SequalPrep Normalization Plate Kit (Invitrogen). Aliquots of the pool were then column (MinElute PCR Purification Kit, Qiagen) and gel purified (QIAquick Gel Extraction Kit, Qiagen). Libraries were then quantified and sequenced with a 600 cycle MiSeq Reagent Kit (270×270; Illumina) with ∼15% PhiX spike-in.

Demultiplexed sequencing reads were processed using QIIME2 v2020.2 (qiime2.org) with denoising by DADA2 [42]. Taxonomy was assigned using the DADA2 implementation of the RDP classifier [43] using the DADA2 formatted training sets for SILVA version 138 (benjjneb.github.io/dada2/assign.html). For Amplicon Sequence Variants (ASV) analysis, we utilized quality scores to set truncation and trim parameters. The reverse read of human 16S data suffered from low sequence quality and reduced the overall ASV counts, so we therefore analyzed only the forward reads, although a separate analysis using merged forward and reverse reads complemented the findings we report in this manuscript. For the manuscript, forward reads were truncated to 220 base pairs and underwent an additional 5 base pairs of trimming for 16S analysis of human stool. For gnotobiotic mice, forward and reverse reads were truncated to 200 and 150 base pairs respectively. ASVs were filtered such that they were present in more than one sample with at least a total of 10 reads across all samples. Alpha diversity metrics were calculated on subsampled reads using Vegan [44] and Picante [45] R packages. The PhILR Euclidean distance was calculated by first carrying out the phylogenetic isometric log ratio transformation (philr, PhILR [46]) followed by calculating the Euclidean distance (vegdist, Vegan [44]). Principal coordinates analysis was carried out using the pcoa function of APE [47]. ADONIS calculations were carried out (adonis, Vegan) with 999 replications on each distance metric. Centered log_2_- ratio (CLR) normalized abundances were calculated using the Make.CLR function in MicrobeR package [48] with count zero multiplicative replacement (zCompositions; [49]). ALDEx2 [50] was used to analyze differential abundances of count data, using features that represented at least 0.05% of total sequencing reads. Corrections for multiple hypotheses using the Benjamini-Hochberg method [51] were performed where applicable. Where described, a false discovery rate (FDR) indicates the Benjamini-Hochberg adjusted *p*-value for an FDR (0.1 unless otherwise specified). Analysis of distance matrices and alpha diversity mirror prior analyses developed in the Turnbaugh lab and were adapted to the current manuscript [9]. Calculations of associations between ASVs and ASA24 questionnaire data were completed by calculating a Spearman rank correlation and then adjusting the *p*-value for a Benjamini-Hochberg FDR using the cor_pmat function in the R package ggcorrplot [52]. The randomForest package [53] was employed to generate random forest classifiers. Given the total number of samples (n=46) we generated 46 classifiers trained on a subset of 45 samples and used each classifier to predict the sample left out. AUCs are visualized utilizing the pROC [54] and ROCR [55] packages.

### Metagenomic sequencing and analysis

Whole-genome shotgun libraries were prepared using the Nextera XT DNA Library Prep Kit (Illumina). Paired ends of all libraries were sequenced on the NovaSeq 6000 platform in a single sequencing run (n=45 subjects; see **Table S2** for relevant metadata and statistics). Illumina reads underwent quality trimming and adaptor removal using fastp [56] and host read removal using BMTagger v1.1.0 (ftp.ncbi.nlm.nih.gov/pub/agarwala/bmtagger/) in the metaWRAP pipeline (github.com/bxlab/metaWRAP) [57]. Metagenomic samples were taxonomically profiled using MetaPhlan2 v2.7.7 [58] and functionally profiled using HUMAnN2 v0.11.2 [59], both with default parameters. Principal coordinates analysis on MetaPhlan2 species-level abundances was carried out using Bray Curtis distances and the pcoa function of APE [47]. Metaphlan2 abundance outputs were converted to counts and subsampled to even sample depth. Differences between groups were determined utilizing the Aldex2 package as described above. Tables of gene family abundances from HUMAnN2 were regrouped to KEGG orthologous groups using humann2_regroup_table. Functional pathways relating to short-chain fatty acid production were manually curated from the pathway outputs from HUMANn2 and normalized by the estimated genome equivalents in each microbial community obtained from MicrobeCensus [60].

### Quantification of bacterial load

Quantitative PCR (qPCR) was performed on DNA extracted from the human stool samples. DNA templates were diluted 1:10 into a 96-well plate. Samples were aliquoted in a 384-well plate, and PCR primers and iTaq Universal Probes Supermix were added utilizing an Opentrons OT-2 instrument then analyzed on a BioRad CFX384 thermocycler with an annealing temperature of 60°C. The following primers including a FAM labeled PCR probe was used for quantification: 891F, TGGAGCATGTGGTTTAATTCGA; 1003R, TGCGGGACTTAACCCAACA; 1002P, [6FAM]CACGAGCTGACGACARCCATGCA[BHQ1]. Absolute quantifications were determined against a standard curve of purified 8F/1542R amplified *Vibrio casei* DNA. Reactions identified as inappropriately amplified by the instrument were rejected, and the mean values were used for downstream analysis. Absolute 16S rRNA gene copy number was derived by adjustments for dilutions during DNA extraction and template normalization dividing by the total fecal mass used for DNA extraction in grams.

### Nuclear magnetic resonance (NMR) metabolomics

NMR spectroscopy was performed at 298K on a Bruker Avance III 600 MHz spectrometer configured with a 5 mm inverse cryogenic probe (Bruker Biospin, Germany) as previously described [61]. Lean and obese EA and W individuals (n=20 total individuals, five in each group) were selected and matched based on body composition and metabolic parameters. Stool samples from these subjects were subjected to NMR-based metabolomics. 50 mg of human feces were extracted with 1 mL of phosphate buffer (K_2_HPO_4_/NaH_2_PO_4_, 0.1 M, pH 7.4, 50% v/v D_2_O) containing 0.005% sodium 3-(trimethylsilyl) [2,2,3,3-2H4] propionate (TSP-d_4_) as a chemical shift reference (δ 0.00). Samples were freeze-thawed three times with liquid nitrogen and water bath for thorough extraction, then homogenized (6500 rpm, 1 cycle, 60 s) and centrifuged (11,180 g, 4 °C, 10 min). The supernatants were transferred to a new 2 mL tube. An additional 600 μL of PBS was added to the pellets, followed by the same extraction procedure described above. Combined fecal extracts were centrifuged (11,180 g, 4°C, 10 min), 600 μL of the supernatant was transferred to a 5 mm NMR tube (Norell, Morganton, NC) for NMR spectroscopy analysis. A standard one-dimensional NOESY pulse sequence noesypr1d (recycle delay-90°-t1-90°-tm-90°-acquisition) is used with a 90 pulse length of approximately 10s (-9.6 dbW) and 64 transients are recorded into 32k data points with a spectral width of 9.6 KHz. NMR spectra were processed as previously described [61]. First, spectra quality was improved with Topspin 3.0 (Bruker Biospin, Germany) for phase and baseline correction and chemical shift calibration. AMIX software (version: 3.9.14, Bruker Biospin, Germany) was used for bucketing (bucket width 0.004 ppm), removal of interfering signal, and scaling (total intensity). Relative concentrations of identified metabolites were obtained by normalized peak area.

### Targeted gas chromatography mass spectrometry (GC-MS) assays

Targeted analysis of short-chain fatty acids (SCFAs) and branched-chain amino acids (BCAAs) was performed with an Agilent 7890A gas chromatograph coupled with an Agilent 5975 mass spectrometer (Agilent Technologies Santa Clara, CA) using a propyl esterification method as previously described [61]. 50 mg of human fecal samples were pre-weighed, mixed with 1 mL of 0.005 M NaOH containing 10 μg/mL caproic acid-6,6,6-d3 (internal standard) and 1.0 mm diameter zirconia/silica beads (BioSpec, Bartlesville, OK). The mixture was thoroughly homogenized and centrifuged (13,200 g, 4°C, 20 min). 500 μL of supernatant was transferred to a 20 mL glass scintillation vial. 500 μL of 1-propanol/pyridine (v/v=3/2) solvent was added into the vial, followed by a slow adding of an aliquot of 100 μL of esterification reagent propyl chloroformate. After a brief vortex of the mixture for 1 min, samples were derivatized at 60°C for 1 hour. After derivatization, samples were extracted with hexane in a two-step procedure (300 μL + 200 μL) as described [62]. First, 300 μL of hexane was added to the sample, briefly vortexed and centrifuged (2,000g, 4°C, 5 min), and 300 μL of the upper layer was transferred to a glass autosampler vial. Second, an additional 200 μL of hexane was added to the sample, vortexed, centrifuged, and the 200 μL upper layer was transferred to the glass autosampler vial. A combination of 500 μL of extracts were obtained for GC-MS analysis. A calibration curve of each SCFA and BCAA was generated with series dilution of the standard for absolute quantitation of the biological concentration of SCFAs and BCAAs in human fecal samples.

### Targeted bile acid quantitation by UHPLC-MS/MS

Bile acid quantitation was performed with an ACQUITY ultra high pressure liquid chromatography (UHPLC) system using a Ethylene Bridged Hybrid C8 column (1,7 µm, 100 mm x 2.1 mm) coupled with a Xevo TQ-S mass spectrometer equipped with an electrospray ionization (ESI) source operating in negative mode (All Waters, Milford, MA) as previously described [63]. Selected ion monitoring (SIM) for non-conjugated bile acids and multiple reaction monitoring (MRM) for conjugated bile acids was used. 50 mg of human fecal sample was pre-weighed, mixed with 1 mL of pre-cooled methanol containing 0.5 μM of stable-isotope-labeled bile acids (internal standards) and 1.0 mm diameter zirconia/silica beads (BioSpec, Bartlesville, OK), followed by thorough homogenization and centrifugation. Supernatant was transferred to an autosampler vial for analysis. 100 µL of serum was extracted by adding 200 µL pre-cooled methanol containing 0.5 μM deuterated bile acids as internal standards. Following centrifugation, the supernatant of the extract was transferred to an autosampler vial for quantitation. Calibration curves of individual bile acids were drafted with bile acid standards for quantitation of the biological abundance of bile acids.

### Gnotobiotic mouse experiments

All mouse experiments were approved by the UCSF Institutional Animal Care and Use Committee and performed accordingly. Germ-free mice were maintained within the UCSF Gnotobiotic Core Facility and fed *ad libitum* autoclaved standard chow diet (Lab Diet 5021). Germ-free adult male C57BL/6J mice between 6-10 weeks of age were used for all the experiments described in this paper. 10 lean subjects in our IDEO cohort were selected as donors for the microbiota transplantation experiments, including 5 EA and 5 W donors. The selected donors for gnotobiotic experiments were matched for phenotypic data to the degree possible (**Table S4**). Stool samples to be used for transplantation were resuspended in 10 volumes (by weight) of brain heart infusion media in an anaerobic Coy chamber. Each diluted sample was vortexed for 1 min and left to settle for 5 min, and a single 200 µL aliquot of the clarified supernatant was administered by oral gavage into each germ-free mouse recipient. In experiments LFPP1 and LFPP2, microbiome transplantations were performed for 2 donors per experiment (1 W, 1 EA) with gnotobiotic mice housed in sterile isolators (CBC flexible, softwall isolator) and maintained on *ad libitum* standard chow also known as low-fat, high-plant-polysaccharide (LFPP) diet. In LFPP1, 6 germ-free mice per colonization group received an aliquot of stool from a donor of either ethnicity and body composition (measured using EchoMRI) were recorded on the day of colonization and at 6 weeks post-transplantation (per group n=6 recipient mice, 1 isolator, 2 cages). In LFPP2, we shortened the colonization time to 3 weeks and used two new donor samples. For the third experiment (HFHS experiment), mice were weaned onto an irradiated high-fat, high-sugar diet (HFHS, TD.88137, Envigo) for four weeks prior to colonization and housed in pairs in Tecniplast IsoCages. The same 4 donors from LFPP1 and LFPP2 were included in the HFHS experiment, in addition to 6 new donors (per donor n=2 recipient mice, 1 IsoCage). Body weight and body composition were recorded on the day of colonization and again at 3 weeks post-transplantation. Mice were maintained on the HFHS diet throughout the experiment. All samples were sequenced in a single pool (**Table S3**). For comparisons between donors and recipient mice, donors and recipient mice were subsampled to even sequencing depth and paired between donor and recipient mice (range: 18,544-78,361 sequencing reads/sample).

### Glucose tolerance tests

Food was removed from mice 10 hr (LFPP1 experiment) or 4 hr (HFHS experiment) prior to assessment of glucose tolerance. Mice received i.p. injections of D-glucose (2 mg/kg), followed by repeated collection of blood by tail nick and determination of glucose levels by handheld glucometer (Abbott Diabetes Care) over a 2-hour period.

### Geographic analyses

Map tiles and distance data was obtained using GGMap [64], OpenStreet Maps [65], and the Imap R [66] packages. GGMap was employed using a Google Cloud API key and the final map tiles were obtained in July 2020 [64]. Spearman ranked correlation coefficients (*rho)* were calculated as embedded in the ggpubr [67] R package. 2018 US Census data for EA and W subjects was obtained (B02001 table for race, data.census.gov) for the ZIP codes available in our study and using the leaflet [68] package. The census data used is included as part of **Table S2** to aid in reproduction. Each census region is plotted as a percentage of W individuals over a denominator of W and EA subjects. The leaflet package utilized ZIP Code Tabulation Areas (ZCTAs) from the 2010 census. We extracted all ZCTAs starting with 9, and the resulting 29 ZIP codes that overlap with IDEO subjects were analyzed (**Table S2)**. Two ZCTAs (95687 and 95401) were primarily W when comparing W and EA subjects. There were two W subjects recruited from these ZTCAs. These ZIP codes are cut off based on the zoom magnification for that figure and as a result ZTCAs for 27 individuals are plotted. Distance to a central point in SF was calculated. The point of reference was latitude=37.7585102, longitude=-122.4539916.

### Dietary questionnaire correlation analysis

DHQIII and ASA24 data were analyzed using a Euclidean distance matrix. These transformations were completed using the cluster package [69]. Subsequent analysis was completed using the vegan package [44]. Procrustes transformations were performed using 16S-seq data from human subjects, which was then subjected to a PhILR transformation. The resulting matrix was rotated against the distance matrix for ASA24 or DHQIII questionnaire data using the procrustes command in the vegan R package using 999 permutations. Mantel statistics were calculated utilizing the mantel command of the vegan package.

### R packages used in this study

Picante [45], PhILR [46], MicrobeR [48], ALDEx2 [50], ggcorrplot [52], randomForest [53], GGMap [64], OpenStreetMap [65], IMap [66], ggpubr [67], leaflet [68], cluster [69], readxl [70], Rtsne [71], vegan [72], ape [73], tigris [74], lmerTest [75], qiime2R [76], gghighlight [77], Phyloseq [78], Janitor [79], table 1 [80], ggplot2 [81].

### Statistical analyses

Statistical analysis of the human data was performed using the table1 package in R (STATCorp LLC. College Station, TX). Human data were presented as mean ± SD. Unpaired independent Student’s *t* tests were used to compare differences between the two groups in the case of continuous data and in the case of categorical data the χ^2^ test was utilized. These tests were adjusted for a Benjamini-Hochberg false discovery rate utilizing the command p.adjust in R, which is indicated as an adjusted *p-*value in the tables. In **Tables S9** and **S10** no values met an adjusted *p-*value cutoff of < 0.1. In **Table S1**, *p*-values indicated by numbers were pooled together for adjustments and those represented by symbols were separately pooled together for adjustment. All microbiome-related analyses were carried out in R version 3.5.3 or 4.0.2. Where indicated, Wilcoxon rank-sum tests were calculated. A Benjamini-Hochberg adjusted *p*-value (FDR) of 0.1 was used as the cutoff for statistical significance unless stated otherwise. Statistical analysis of glucose tolerance tests was carried out using linear mixed effects models with the lmerTest [75] R package and mouse as random effect. Graphical representation was carried out using ggplot2. Boxplots indicate the interquartile range (25th to 75th percentiles), with the center line indicating the median and whiskers representing 1.5x the interquartile range.

## RESULTS

Ethnicity was associated with marked differences in the human gut microbiota. Principal coordinates analysis of PhILR Euclidean distances from 16S-seq data (**Table S2,** n=22 EA, 24 W subjects) revealed a subtle but significant separation between the gut microbiotas of EA and W subjects (*p*=0.007, R^2^=0.046, ADONIS; **Fig. 1A**). Statistical significance was robust to the distance metric used (**Table S5**). Bacterial diversity was significantly higher in W individuals across three distinct metrics: Faith’s phylogenetic diversity, ASV richness, and Shannon diversity (**Fig. 1B**). Six bacterial phyla were significantly different between ethnicities (**Fig. 1C**), with only one phylum, *Verrucomicrobiota*, enriched in W subjects. Significant differences were also detectable at the genus level (**Fig. 1D**). We identified two ASVs that were significantly different between ethnicities: *Blautia obeum* and a *Streptococcus* species, both enriched in EA subjects **(Fig. 1E)**. There were no significant differences between ethnicities in overall colonization level (**Fig. 1F**).

**Figure 1.**
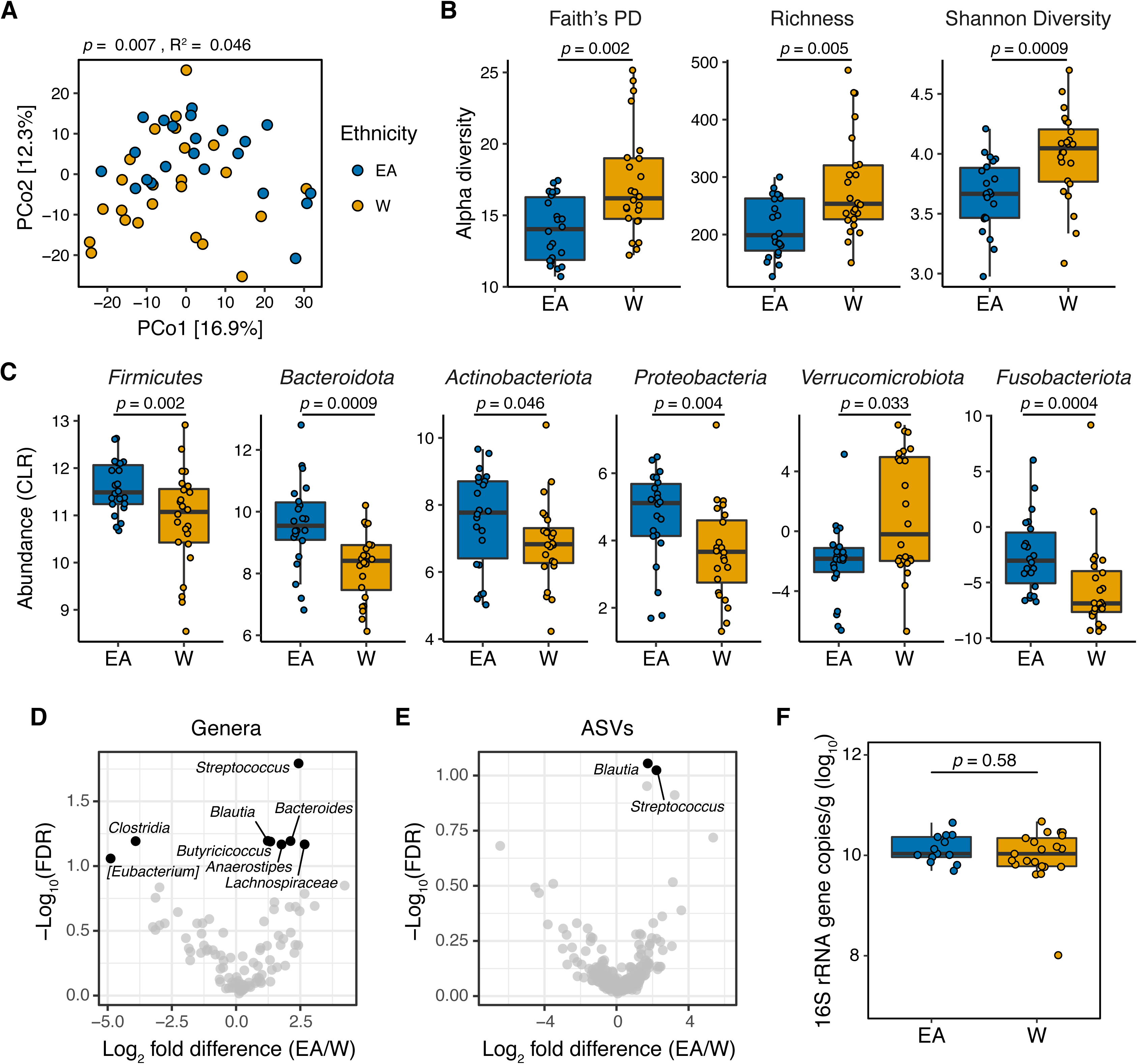
The gut microbiota is distinct between East Asian and White subjects living in the Bay Area. **(A-C)** Each point represents a single individual’s gut microbiota based upon 16S-seq. **(A)** Principal coordinate analysis of PhILR Euclidean distances reveals significant separation between ethnic groups (ADONIS test values shown). Additional distance calculations for complementary distance matrix calculations are shown in **Table S5**. **(B)** Calculations of alpha diversity between EA and W subjects. *p*-values determined using Wilcoxon rank-sum tests. **(C)** Six bacterial phyla are significantly different between EA and W individuals (*p*<0.05, Wilcoxon rank-sum test). **(D,E)** Volcano plot of ALDEx2 differential abundance testing on **(D)** genera and **(E)** ASVs detected by 16S-seq in the gut microbiotas of EA versus W individuals. Significantly different (FDR < 0.1) features are highlighted in black and labeled by genus or the most specific taxonomic assignment. **(A-E)** n=22 EA and n=24 W individuals. **(F)** No significant difference in overall gut microbial colonization assessed by qPCR quantification of 16S rRNA gene copies per gram wet weight (n=13 EA, n=21 W, Wilcoxon rank-sum test).

Phylogenetic analyses of all ASVs revealed marked variations in the direction of change across different phyla between EA and W subjects (**Fig. S1A**), indicating that the phylum level trends (**Fig. 1C**) resulted from the integration of subtle shifts across multiple component members. Next, we used a random forest classifier to define biomarkers in the gut microbiota that distinguish EA and W subjects (**Figs. S1B-D**). Classifiers employing ASV data and PhILR transformed phylogenetic nodes were trained using leave-one-out cross-validation. *Blautia obeum* (ASV1) was the top contributor to the resulting classifier, followed by *Anaerostipes hadrus* (ASV45) and then *Streptococcus parasanguinis* (ASV110) (**Fig. S1B, Table S6)**. Both classifiers demonstrated the ability to distinguish between ethnic groups, with PhILR transformed phylogenetic nodes achieving a higher area under the curve compared to ASVs (**Figs. S1C,D**).

Metagenomic sequencing provided independent confirmation of differences in the gut microbiome between ethnicities (**Table S2,** n=21 EA, 24 W subjects). Consistent with our 16S- seq analysis, we detected a subtle but significant difference in the gut microbiomes between ethnicities based upon metagenomic species abundances (*p*=0.004, R^2^=0.047, ADONIS) and gene families (*p*=0.029, R^2^=0.036, ADONIS). Visualization of species within each phylum revealed marked variation in the magnitude and direction of change between ethnicities (**Fig. 2A**). Four bacterial species were found to be significantly different between ethnicities in our metagenomic data: W individuals had higher levels of *Akkermansia muciniphila*, *Bacteroidales bacterium* ph8, and *Roseburia hominis,* and lower levels of *Ruminococcus gnavus,* compared to EA individuals (**Fig. 2B**).

**Figure 2.**
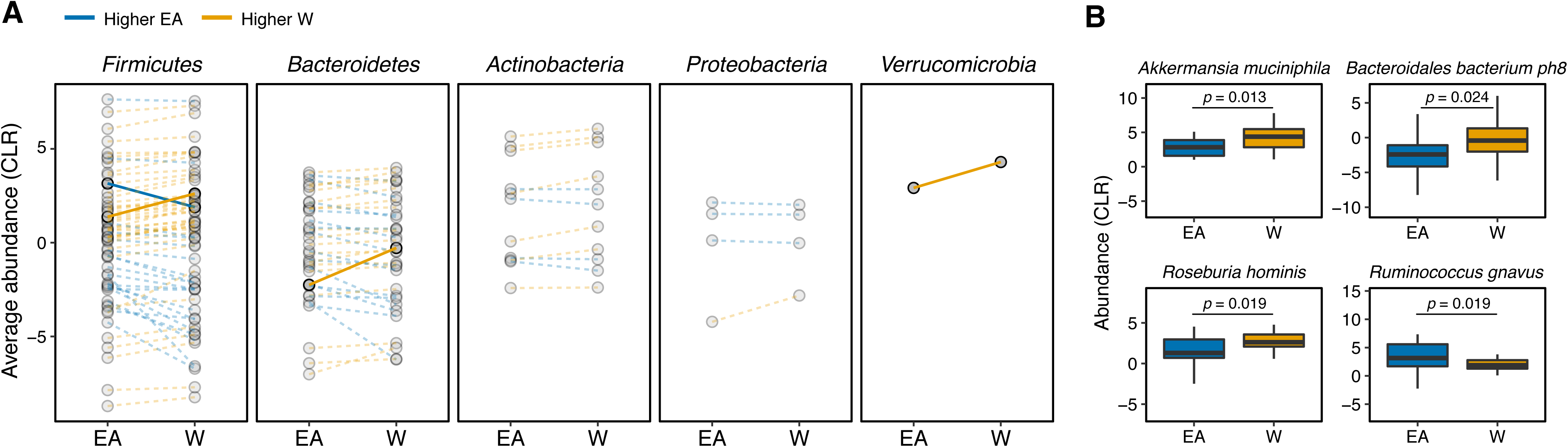
Metagenomic sequencing corroborates differences in the gut microbiota between ethnicities. **(A)** Each point represents the average relative abundance for a given species within each ethnic group, connected with a line that is colored by the ethnic group with higher mean abundance of each species: EA (blue) and W (orange). Solid lines highlight four bacterial species that are significantly different between ethnicity (*p*<0.05, ALDEx2). **(B)** Abundances of the four species found to be significantly different between ethnicities. *p*-values determined using Wilcoxon rank-sum tests for individual plots. **(A,B)** n=21 EA and n=24 W individuals.

Next, we used NMR-based stool metabolomics to gain insight into the potential functional consequences of ethnicity-associated differences in the human gut microbiome (**Table S2**, n=10 subjects/ethnicity). Metabolite profiles were more strongly associated with ethnicity (*p*=0.008, R^2^=0.128, ADONIS; **Fig. 3A**) than community structure (R^2^=0.029-0.055, ADONIS; **Table S5**) or gene abundance (*p*=0.029, R^2^=0.036, ADONIS). Feature annotations revealed elevated levels of the branched chain amino acid (BCAA) valine and the short-chain fatty acids (SCFAs) acetate and propionate in EA subjects (**Fig. 3B** and **Table S7**). In contrast, proline, formate, alanine, xanthine, and hypoxanthine were found at higher levels in W subjects (**Fig. 3B**). To assess the statistical significance and reproducibility of these trends, we used targeted GC-MS and UPLC-MS/MS to quantify a panel of BCAAs, SCFAs, and bile acids (**Table S8**). Confirming our NMR data, EA subjects had significantly higher levels of stool acetate (**Fig. 3C**) and propionate (**Fig. 3D**); however, we did not detect any significant differences in BCAAs or bile acids **(Fig. S2**). Isobutyrate (which was not detected by NMR) was also significantly higher in EA subjects (**Fig. 3E**). In agreement with these metabolite levels, a targeted re-analysis of our metagenomic data revealed a significant enrichment in two SCFA-related pathways: “pyruvate fermentation to butanoate” (*p*=0.023, fold-difference=2.216) and “superpathway of *Clostridium acetobutylicum* acidogenic fermentation” (*p*=0.023, fold-difference=2.182).

**Figure 3.**
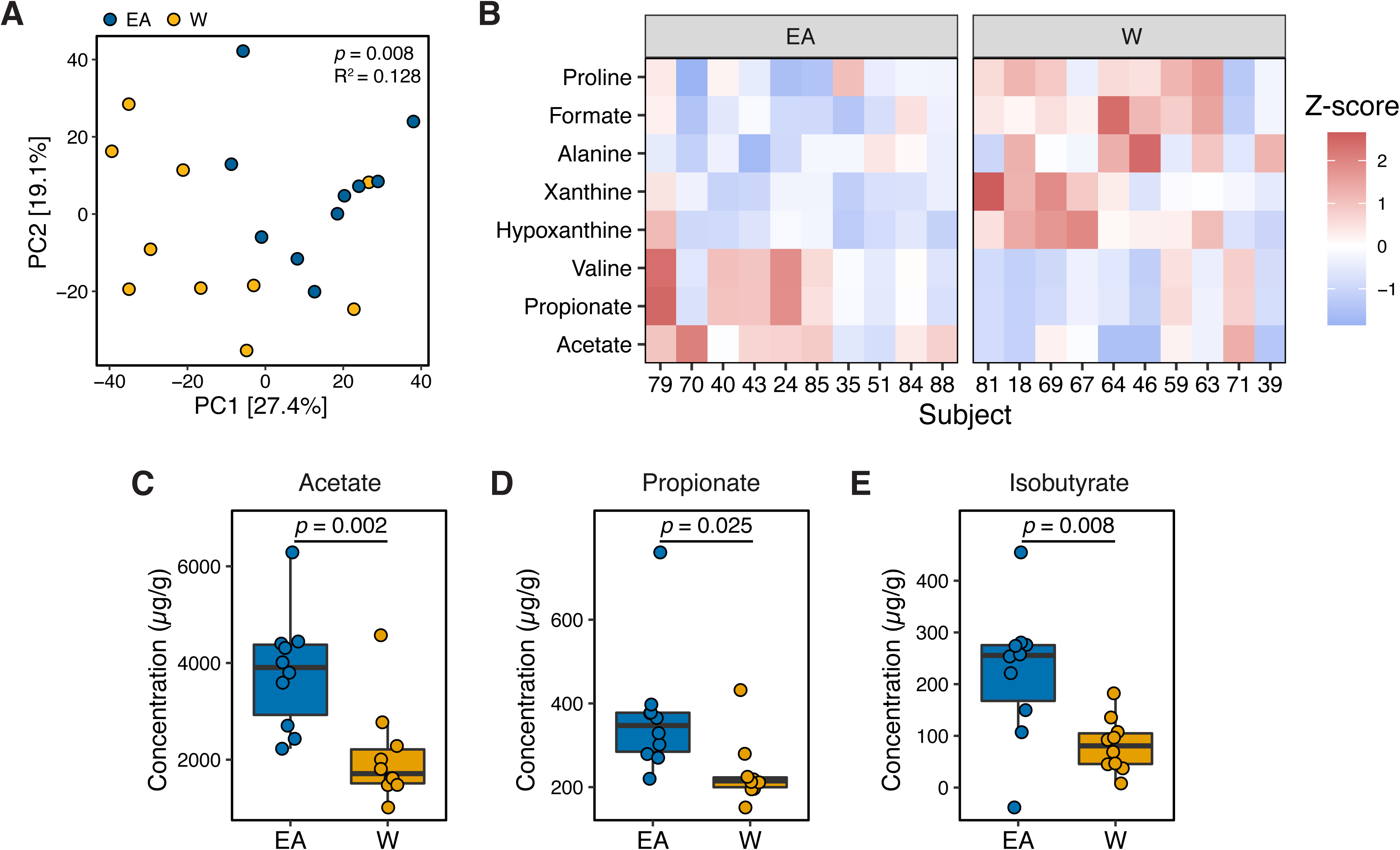
Metabolomics and targeted metabolite profiling highlight significant differences in bacterial fermentation end-products between ethnicities. **(A)** Global profiling of the stool metabolome by proton nuclear magnetic resonance (^1^H NMR) revealed a significant separation in metabolomic profiles between EA and W individuals (ADONIS test values shown). **(B)** Representative stool metabolites contributing to the separation of stool metabolomic profiles between EA and W individuals (*p*<0.05, Wilcoxon rank-sum test). **(C-E)** Gas chromatography-mass spectrometry analysis of short-chain fatty acids (SCFAs) revealed significantly higher concentrations of acetate **(C)**, propionate **(D)** and isobutyrate **(E)** in the stool samples of EA compared to W individuals. *p*-values determined using Wilcoxon rank-sum tests. **(A-E)** n=10 EA and n=10 W individuals.

Consistent with prior work [23, 82], we found that gut bacterial richness in W individuals was significantly associated with both BMI (**Fig. 4A**) and body fat percentage (**Fig. 4B**). Remarkably, these associations were undetectable in EA subjects (**Figs. 4A,B**) even when other metrics of bacterial diversity were used (**Fig. S3**), with the single exception of a negative correlation between Shannon diversity and BMI in EA subjects **(Fig. S3C)**. Re-analysis of our data separating lean and obese individuals revealed that the previously observed differences between ethnic groups were driven by lean individuals. Compared to lean EA individuals, lean W subjects had significantly higher bacterial diversity (**Fig. 4C**), in addition to greater differences in both gut microbial community structure (*p*=0.0003, R^2^=0.122, ADONIS; **Fig. 4D**) and metabolite profiles (*p*=0.010, R^2^=0.293, ADONIS; **Fig. 4E**). By contrast, obese W versus EA individuals were not different across any of these metrics (**Figs. 4C-E**), except for lower Shannon diversity in obese EA compared to W individuals (**Fig. 4C**). We also detected differences in the gut microbiotas of lean EA and W individuals at the phylum (**Fig. 5A**) and genus (**Fig. 5B**) levels that were largely consistent with our original analysis of the full dataset (**Figs. 1C,D**). More modest differences in the gut microbiota between ethnicities were observed in obese subjects (**Figs. 5A,C**).

**Figure 4.**
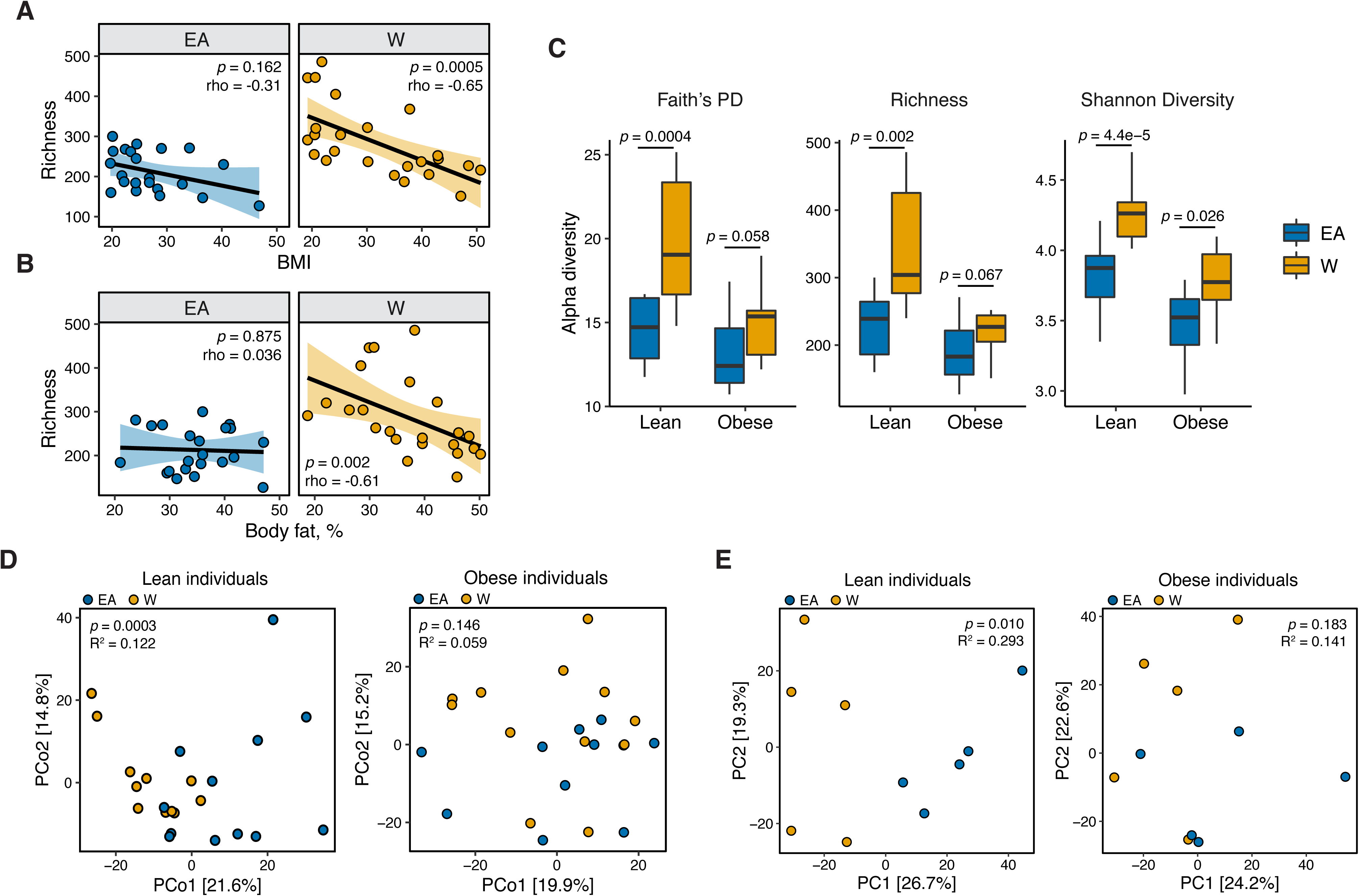
Ethnicity-associated differences in gut microbial diversity and community structure are more pronounced in lean individuals. **(A,B)** Bacterial richness is negatively correlated with (A) BMI and (B) percent body fat in W but not EA individuals (Spearman rank correlation coefficients and *p-*values are shown for each graph). **(C)** Microbial diversity metrics are more distinct between ethnic groups in lean relative to obese individuals. *p*-values determined using Wilcoxon rank-sum tests. **(D)** Principal coordinate analysis of PhILR Euclidean distances reveals significant separation between the gut microbiotas of EA and W lean individuals, with no separation in obese subjects (ADONIS test values shown). **(A-D)** n=12 EA lean, 10 EA obese, 11 W lean, and 13 W obese individuals. **(E)** Global profiling of the stool metabolome by proton nuclear magnetic resonance (^1^H NMR) stratified by lean and obese individuals reveals a significant difference in the metabolomic profiles of lean EA and W individuals that is not detectable in obese individuals (ADONIS test values shown; n=5 individuals/group).

**Figure 5.**
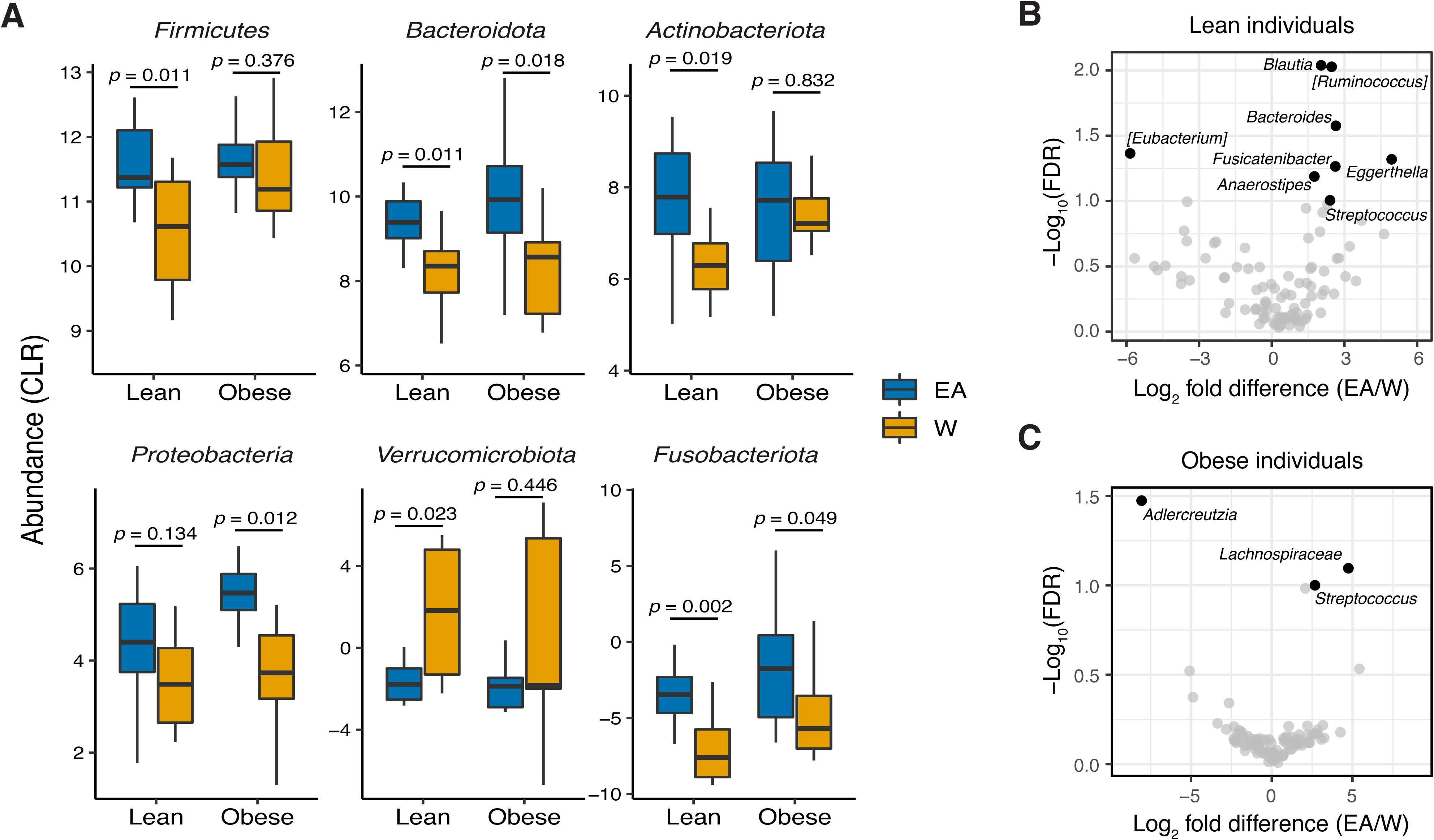
Ethnicity-associated bacterial taxa in lean and obese individuals. **(A)** 5/6 phyla differentially abundant between ethnicities (see Fig. 1C) were also significantly different between lean EA and W individuals. Three phyla were significantly different between obese EA and W individuals (*p*<0.05, Wilcoxon rank-sum test). **(B,C)** Volcano plot of ALDEx2 differential abundance testing on genera in stool microbiotas of lean **(B)** and obese **(C)** EA versus W individuals, with significantly different genera highlighted (FDR<0.1, ALDEx2). **(A-C)** n=12 EA lean, 10 EA obese, 11 W lean, and 13 W obese individuals.

Next, we sought to understand the potential drivers of differences in the gut microbiome between ethnic groups in lean individuals within the IDEO cohort. Consistent with prior studies [83], PERMANOVA analysis of our full 16S-seq dataset revealed that diabetes [84], age [85], metformin use [86], and statin intake [87] to be significantly associated with variance in the PhILR Euclidean distances **(Fig. S4)**; however, none of these factors were significantly different between ethnicities (**Table S1**). Although everyone in the cohort was recruited from the San Francisco Bay Area, birth location varied widely (**Fig. S5**). There was no significant difference in the proportion of subjects born in the USA between ethnicities (75% W, 54.5% EA; *p*=0.15, Pearson’s χ^2^ test). There was also no significant difference in the geographical distance between birth location and San Francisco [W median 2,318 (2.2-6,906) miles; EA median 1,986 (2.2-6,906) miles; *p*=0.69, Wilcoxon rank-sum test) or the amount of time spent in the San Francisco Bay Area at the time of sampling [W median 270 (8.00-741) months; EA median 282.5 (8.50-777) years; *p*=0.42, Wilcoxon rank-sum test). While obese subjects were markedly distinct from lean individuals of both ethnicities with regard to measured metabolic and laboratory parameters (**Table S1**), there were no statistically significant differences between ethnic groups after separating lean and obese individuals (**Table S1**).

Surprisingly, we did not detect any significant differences in either short- (**Table S9**) or long-term (**Table S10**) dietary intake between ethnicities. Consistent with this, Procrustes analysis did not reveal any significant associations between dietary intake and gut microbial community structure: procrustes *p=*0.280 (DHQIII) and *p*=0.080 (ASA24) relative to PhILR transformed 16S- seq ASV data. The Spearman Mantel statistic was also non-significant [*r*=0.0524, *p*=0.243 (DHQIII) and *r=-*0.0173, *p*=0.590 (ASA24)], relative to PhILR transformed 16S-seq ASV data. Despite the lack of an overall association between reported dietary intake and the gut microbiota, we were able to identify 6 ASVs associated with dietary intake in lean W individuals (**Fig. S6**). In contrast, there was only one significant ASV-level association in lean EA subjects.

Given the marked variation in the gut microbiome at the continental scale [1–3], we hypothesized that the observed differences in lean EA and W individuals may be influenced by a participant’s current address at the time of sampling. Consistent with this hypothesis, we found clear trends in ethnic group composition across ZIP codes in the IDEO cohort (**Figs. 6A,B**) that were mirrored by the 2018 US census data (Pearson *r*=0.52, *p*=0.026 for neighborhoods with greater than 50% white subjects; **Fig. 6D**). Obese individuals from both ethnicities and lean W subjects tended to live closer to the center of San Francisco relative to lean EA subjects (**Fig. 6C**). Distance between current ZIP code and the center of San Francisco and duration of residency within San Francisco were both associated with gut microbial community structure (**Figs. 6E,F**). The association between current address and the gut microbiota was robust to the central point used, as evidenced by using the Bay Bridge as the central reference point (*p*=0.008, rho=0.394, Spearman correlation).

**Figure 6.**
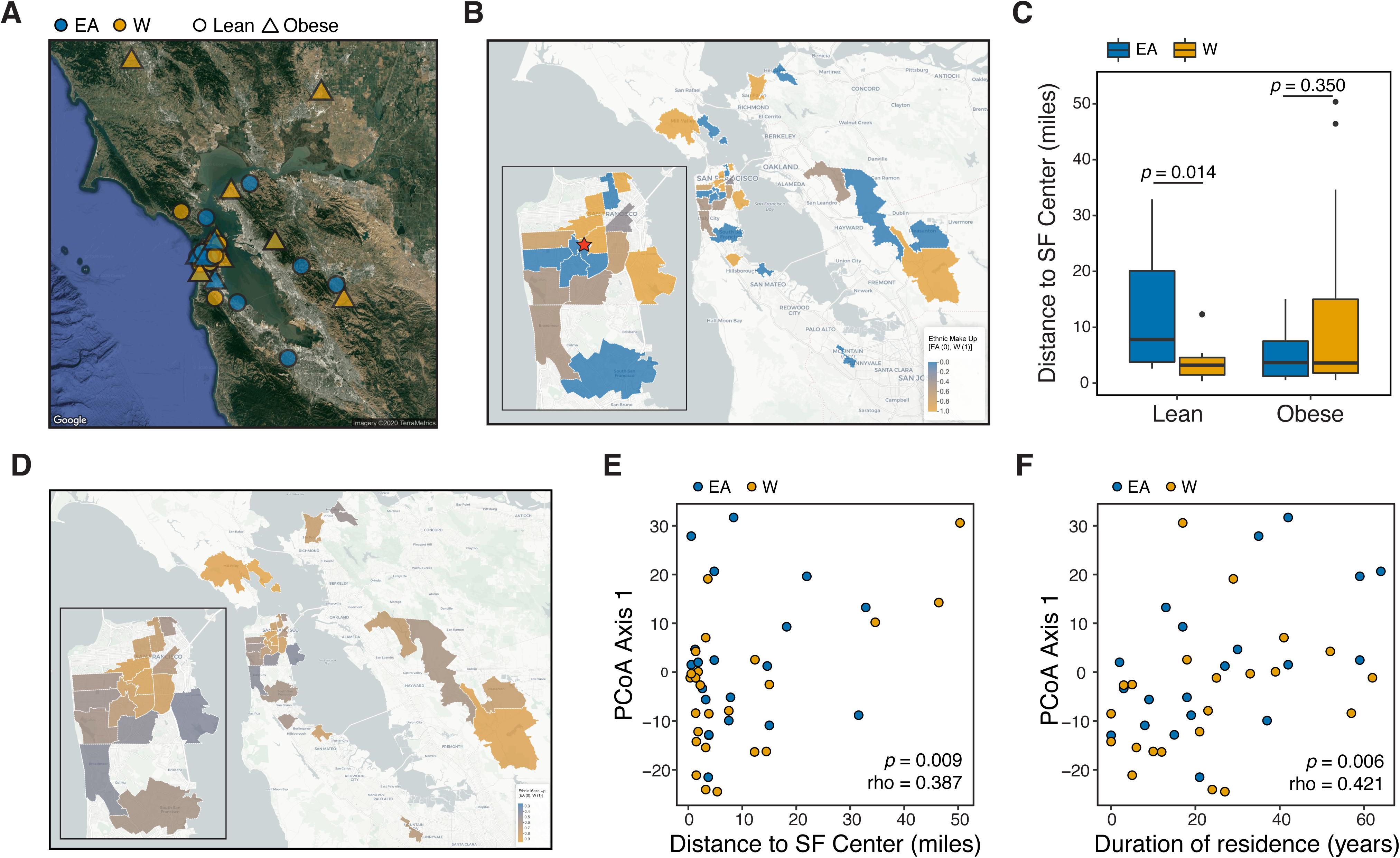
Ethnicity-associated differences in the gut microbiota of lean individuals correlate with geographic location. **(A)** Each symbol represents a subject’s ZIP code. Symbols are colored by ethnicity with shape representing lean and obese subjects (n=44, data was unavailable for 2 subjects; **Table S2**). **(B)** A subset of ZIP Code Tabulation Areas (ZCTAs) zoomed in to focus on San Francisco are colored by the proportion of each ethnicity (n=27 ZTCAs). The red star indicates a central point (latitude=37.7585102, longitude=-122.4539916) within San Francisco used for distances calculated in (C). **(C)** Distance to the center of San Francisco, which is indicated by a star in (B), for IDEO subjects stratified by ethnicity and BMI (n=9-13 individuals/group, *p*-values indicate Wilcoxon rank-sum test). **(D)** US census data for EA and W residents in ZCTAs from (B) is displayed by ethnic make-up (a total of 489,117 W and 347,200 Asian individuals in these areas). **(E,F)** PCoA principal coordinate axis 1 from PhILR Euclidean distances of the 16S-Seq data is significantly correlated with **(E)** the distance of subject’s ZIP code to the center of San Francisco and **(F)** the subject’s duration of residence in the SF Bay Area (n=44 subjects; Spearman rank correlation).

Taken together, our results support the hypothesis that there are stable ethnicity-associated signatures within the gut microbiota of lean EA vs. W individuals that are independent of diet. To experimentally test this hypothesis, we transplanted the gut microbiota of a representative lean W and lean EA individual into germ-free C57BL/6J mice fed a low-fat, high-plant-polysaccharide (LFPP) diet in two separate experiments (per group n=12 mice, 2 donors; per donor n=6 mice, 1 isolator; **Fig. S7A,B**). The donors for this and the subsequent experiments were matched for their metabolic and other phenotypes to minimize potential confounding factors (**Table S4**). Despite maintaining the genetically identical recipient mice on the same autoclaved LFPP diet, we detected significant differences in gut microbial community structure (**Fig. 7A**), bacterial richness (**Fig. 7B**), and taxonomic abundance (**Figs. 7C,D**) between the two ethnicity-specific recipient groups, and these remarkably reflected the differences we saw when initially analyzing the stool samples of the human participants themselves. Notably, we observed significantly lower bacterial richness **(Fig. 7B)** and higher abundance of *Bacteroides* **(Fig. 7D)** in recipient mice transplanted with microbiota from EA compared to W donors.

**Figure 7.**
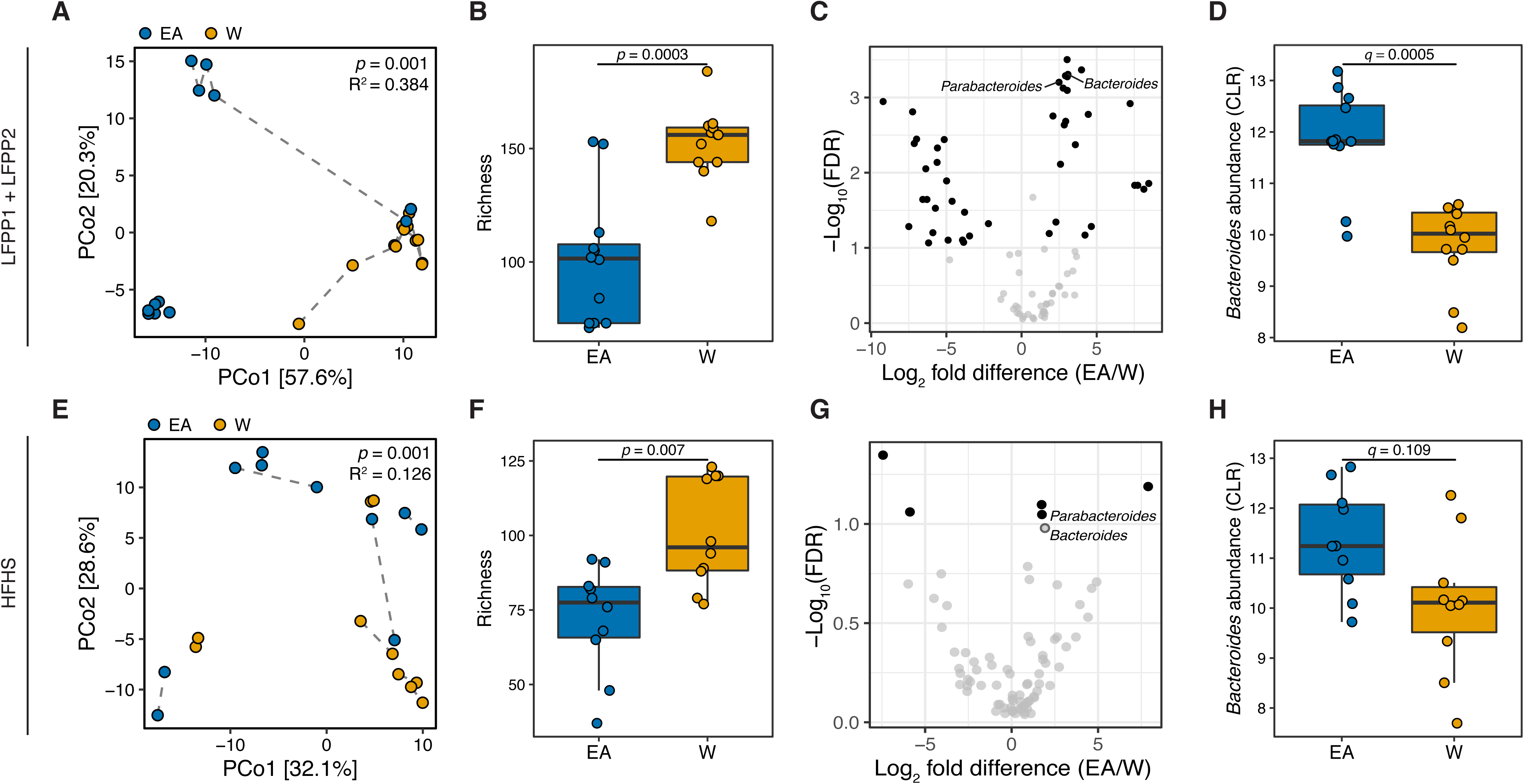
Differences in the human gut microbiota between ethnicities are maintained following transplantation to germ-free mice. **(A,E)** Principal coordinate analysis of PhILR Euclidean distances of stool from germ-free recipient mice transplanted with stool microbial communities from lean EA or W donors and fed either a LFPP (**A**, combined results from two independent experiments; n=12 recipient mice per group) or HFHS (**E**, n=10 recipient mice per group) diet. Significance was assessed by ADONIS. Germ-free mice receiving the same donor sample are connected by a dashed line. Experimental designs are shown in **Fig. S7**. **(B,F)** Bacterial richness is significantly higher in mice who received stool samples from lean W donors compared to EA donors on both the LFPP **(B)** and HFHS **(F)** diets. *p*-values determined using Wilcoxon rank-sum tests. **(C,G)** Volcano plot of ALDEx2 differential abundance testing on genera in the stool microbiomes between transplant groups. The x-axis represents the fold difference between EA (numerator) and W (denominator) subjects. The y-axis is proportional to the false discovery rate (FDR). Black dots indicate significantly different genera (FDR<0.1). *Bacteroides* and *Parabacteroides* (labelled in the volcano plots) are more abundant in mice that received stool samples from EA compared to W donors on both the LFPP **(C)** and HFHS **(G)** diets. **(D,H)** Abundance of the *Bacteroides* genus in mice fed the LFPP **(D)** and HFHS **(H)** diets (ALDEx2 FDR shown).

To assess the reproducibility of these findings across multiple donors and in the context of a distinctive dietary pressure, we next fed 20 germ-free mice a high-fat, high-sugar (HFHS) diet for 4 weeks prior to colonization with microbiota from 5 different W vs. EA donors, and then maintained the mice on this diet following colonization (per group n=10 mice, 5 donors; per donor n=2 mice, 1 cage; **Fig. S7C**). This experiment replicated our original findings on the LFPP diet, including significantly altered gut microbial community structure (**Fig. 7E**), significantly increased richness in mice receiving W donor microbiota (**Fig. 7F**), and a trend towards higher levels of *Bacteroides* in mice receiving the gut microbiotas of EA donors (**Figs. 7G,H**). Of note, the variance explained by ethnicity was lower in mice fed the HFHS diet (R^2^=0.126) than the LFPP diet (R^2^=0.384), potentially suggesting that in the context of human obesity, chronically excessive fat and sugar consumption may serve to diminish the signal otherwise associated with ethnicity. As expected [88–90], the input donor microbiota was distinct from that of the recipient mice (**Fig. S8A**); however, there was no difference between ethnic groups in the efficiency of engraftment (**Figs. S8B-D**). Also consistent with prior data [88], an analysis of the combined data from LFPP and HFHS fed mice revealed marked differences in the gut microbiota between the two diets despite persistent differences between ethnicities (**Fig. S9**).

Finally, mice transplanted with gut microbiomes of EA and W individuals displayed differences in body composition. Mice that received W donor microbiota and were fed the LFPP diet significantly increased their adiposity in conjunction with a reduction in lean mass, relative to mice that received the EA donor microbiota (**Figs. 8A-C**). These trends were mirrored in the human microbiota recipient mice that were fed the HFHS diet **(Figs. 8E-G)**, though they did not reach statistical significance, suggesting that dietary input can mask the metabolic consequences of ethnicity-associated differences in the gut microbiota. There were no significant differences in glucose tolerance between mice receiving stool transplants from donors of different ethnicity in either experiment (**Figs. 8D,H**).

**Figure 8.**
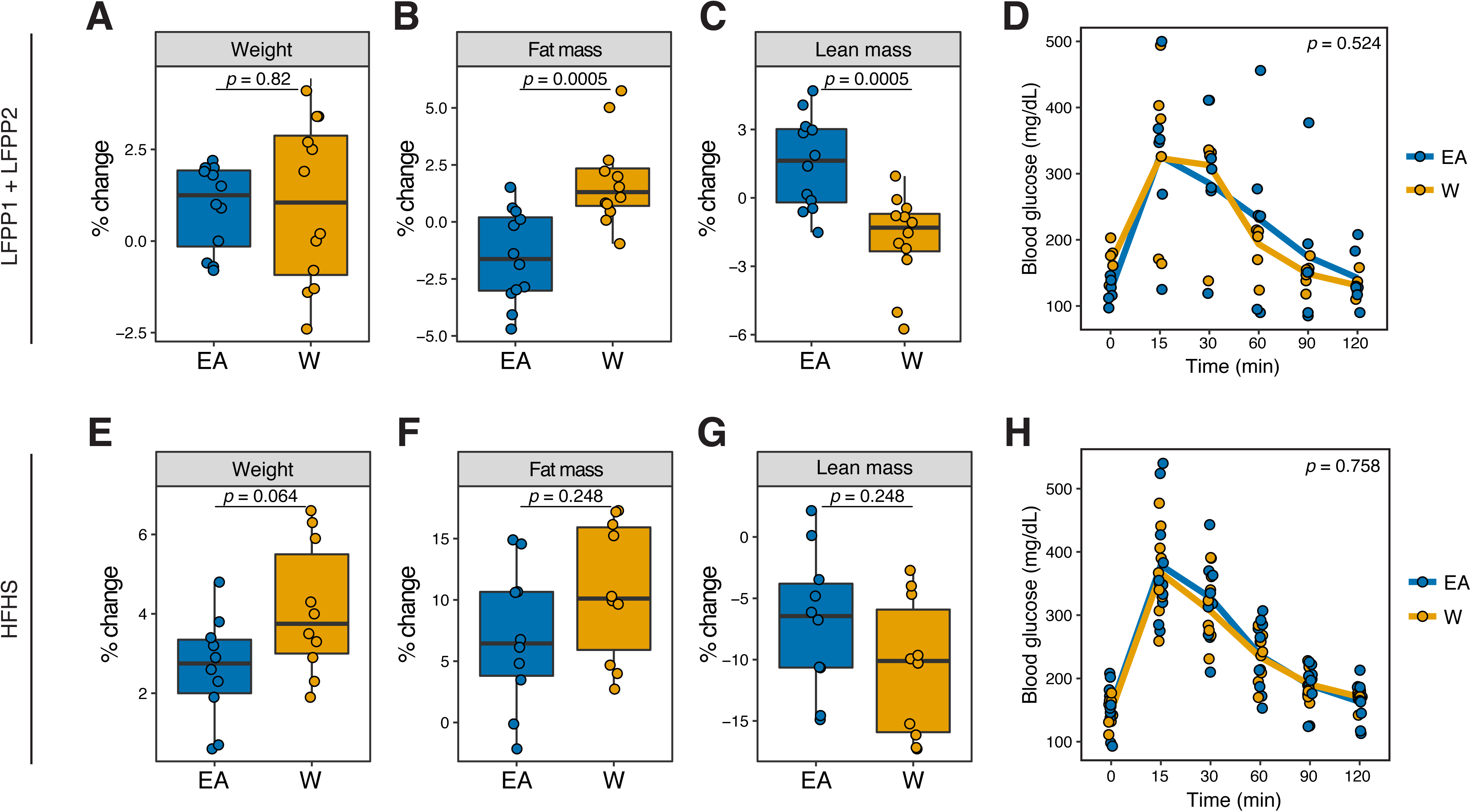
Microbiome transplantation of samples from EA and W individuals differentially affects the body composition of genetically identical recipient mice. **(A-C, E-G)** Percent change in body weight **(A,E)**, fat mass **(B,F)**, and lean mass **(C,G)** relative to baseline are shown on the LFPP (**A-C**) and HFHS (**E-G**) diets. *p*-values determined using Wilcoxon rank-sum tests. **(D,H)** Glucose tolerance test results were not significantly different between groups in either experiment. *p*-values determined using linear mixed effects models with mouse as a random effect. **(A-C)** n=12 recipient mice per group (combined data from two independent experiments). **(D)** n=6 recipient mice per group from a single experiment. **(E-H)** n=10 recipient mice per group. Experimental designs are shown in **Fig. S7** and donor phenotypic data is in **Table S4**.

## DISCUSSION

Despite the potential for immigration to erase some of the geographically specific aspects of gut microbiome structure [3], our study suggests that even in a given geographic location, there remain stable long-lasting microbial signatures of ethnicity, as revealed here for W and EA residents of the San Francisco Bay Area. The mechanisms responsible remain to be elucidated. In lean individuals within the IDEO cohort, these differences appear to be independent of immigration status, host phenotype, or dietary intake. Our experiments using inbred germ-free mice support the stability of ethnicity-associated differences in the gut microbiota on both the LFPP and HFHS diets, while also demonstrating that variations in host genetics are not necessary to maintain these signatures, at least over short timescales.

Our data also supports a potential role for geographic location of residence in reinforcing differences in the gut microbiota between ethnic groups. The specific reasons why current location matters remain unclear. It may reflect subtle differences in dietary intake (*e.g.,* ethnic foods, food sources, or phytochemical contents) that are hard to capture using the validated nutritional surveys employed here [91]. Alternative hypotheses include biogeographical patterns in microbial dispersion [92] or a role for socioeconomic factors, which are correlated with neighborhood [93].

Surprisingly, our findings demonstrate that ethnicity-associated differences in the gut microbiota are stronger in lean individuals. Obese individuals did not exhibit as clear a difference in the gut microbiota between ethnic groups, either suggesting that established obesity or its associated dietary patterns can overwrite long-lasting microbial signatures or alternatively that there is a shared ethnicity-independent microbiome type that predisposes individuals to obesity. Studies in other disease areas (*e.g.,* inflammatory bowel disease and cancer) with similar multi-ethnic cohorts are essential to test the generalizability of these findings and to generate hypotheses as to their mechanistic underpinnings.

Our results in humans and mouse models support the broad potential for downstream consequences of ethnicity-associated differences in the gut microbiome for metabolic syndrome and potentially other disease areas. However, the causal relationships and how they can be understood in the context of the broader differences in host phenotype between ethnicities require further study. While these data are consistent with our general hypothesis that ethnicity-associated differences in the gut microbiome are a source of differences in host metabolic disease risk, we were surprised by both the nature of the microbiome shifts and their directionality. Based upon observations in the IDEO [18] and other cohorts [15, 16], we anticipated that the gut microbiomes of lean EA individuals would promote obesity or other features of metabolic syndrome. In humans, we did find multiple signals that have been previously linked to obesity and its associated metabolic diseases in EA individuals, including increased Firmicutes [9, 94], decreased *A. muciniphila* [95, 96], decreased diversity [23], and increased acetate [97, 98]. Yet EA subjects also had higher levels of *Bacteroidota* and *Bacteroides*, which have been linked to improved metabolic health [99]. More importantly, our microbiome transplantations demonstrated that the recipients of the lean EA gut microbiome had less body fat despite consuming the same diet. These seemingly contradictory findings may suggest that the recipient mice lost some of the microbial features of ethnicity relevant to host metabolic disease or alternatively that the microbiome acts in a beneficial manner to counteract other ethnicity-associated factors driving disease.

EA subjects also had elevated levels of the short chain fatty acids propionate and isobutyrate. The consequences of elevated intestinal propionate levels are unclear given the seemingly conflicting evidence in the literature that propionate may either exacerbate [100] or protect [101] from aspects of metabolic syndrome. Clinical data suggests that circulating propionate may be more relevant for disease than fecal levels [102], emphasizing the importance of considering both the specific microbial metabolites produced, their intestinal absorption, and their distribution throughout the body. Isobutyrate is even less well-characterized, with prior links to dietary intake [103] but no association with obesity [104].

There are multiple limitations of this study. Due to the investment of resources into ensuring a high level of phenotypic information on each cohort member, and due to its restricted geographical catchment area, the IDEO cohort was relatively small at the time of this analysis (n=46 individuals). This study only focused on two of the major ethnicities in the San Francisco Bay Area; as IDEO continues to expand and diversify its membership, we hope to study a sufficient number of participants from other ethnic groups in the future. Stool samples were collected at a single time point and analyzed in a cross-sectional manner. While we used validated tools from the field of nutrition to monitor dietary intake, we cannot fully exclude subtle dietary differences between ethnicities [105], which would be possible with a controlled feeding study [94]. In our animal studies, we focused on metabolically healthy wild-type germ-free mice, only compared two donors per ethnicity in mice fed the LFPP, and did not directly compare the same inocula administered to mice on different diets. Follow-up studies in the appropriate disease models coupled to controlled experimentation with individual strains or more complex synthetic communities are necessary to elucidate the mechanisms responsible for ethnicity-associated changes in host physiology and their relevance to disease.

## CONCLUSIONS

Our results support the utility of considering ethnicity as a covariate in microbiome studies, due to the ability to detect signals that are difficult to capture by more specific metadata such as individual dietary intake values. On the other hand, these findings raise the importance of dissecting the sociological and biological components of ethnicity with the goal of identifying factors that shape the gut microbiota, either alone or in combination. This emerging area of microbiome research is just one component in the broader efforts to explore the boundaries and mechanistic underpinning of ethnicity with respect to multiple ethnic groups. The IDEO cohort provides a valuable research tool to conduct prospective longitudinal and intervention studies examining diabetes including Asian American participants. More broadly, IDEO provides a framework to approach other disease states where self-identified race or ethnicity are thought to contribute to health outcomes related to the microbiome, including the use of gnotobiotic mouse models to examine the specific role of microbial communities in contributing to phenotypes linked to ethnicity. By understanding the biologic features that drive differences between ethnic groups, we may be able to achieve similar health outcomes and to support more precise therapies informed by a broader appreciation of both microbial and human diversity.

## FIGURE LEGENDS

**Figure S1.**
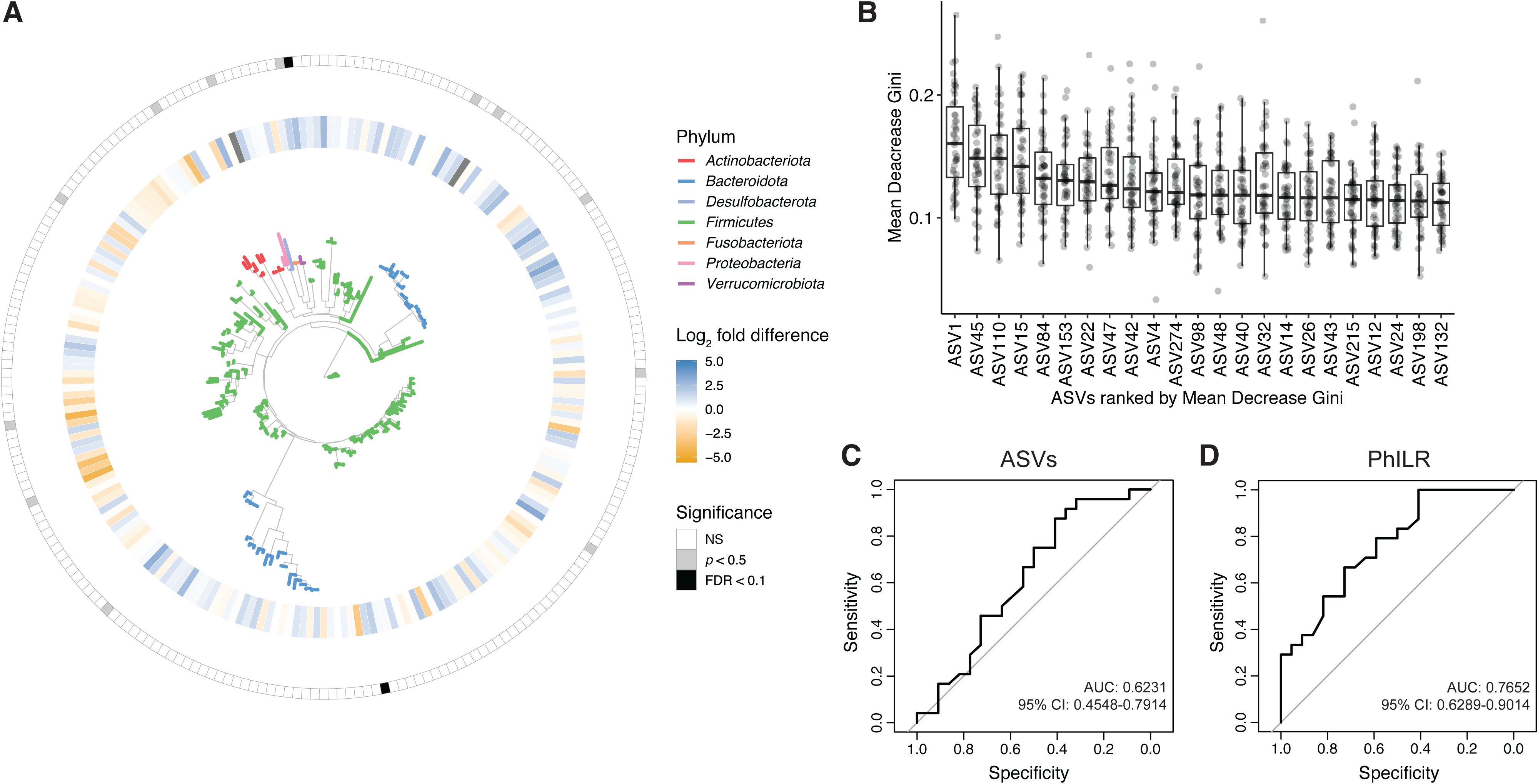
Prediction of ethnicity utilizing Random Forest Classifier from ASV and PhILR transformed phylogenetic nodes. **(A)** A phylogenetic tree of all ASVs generated from 16S-Seq is shown. Leaves are colored by phyla. The inner circle indicates differential abundance (EA_CLR_- W_CLR_) between ethnicities. The outer circle is colored by whether a given ASV was statistically significant by Welch’s *t*-test (gray) or both a Welch’s *t*-test and FDR < 0.1 (black bars; labeled in Fig. 1E). **(B-D)** A Random Forest classifier was developed utilizing ASV data **(B,C)** and PhILR transformed ASV data (**D**) representing phylogenetic nodes on the tree visualized in panel A. 46 classifiers were trained on a subset of 45 individuals and then used to predict the remaining individual (leave-one-out cross-validation). **(B)** ASVs in the top 90th percentile for median mean decrease Gini are plotted. Each dot represents the value for mean decrease Gini for a given classifier (n=46 total classifiers made up of a subset of 45 samples). **(C-D)** AUCs of classifiers for ASV data **(C)** and phylogenetic nodes obtained utilizing PhILR transformation **(D)** are plotted with values of AUC and 95% confidence intervals displayed.

**Figure S2.**
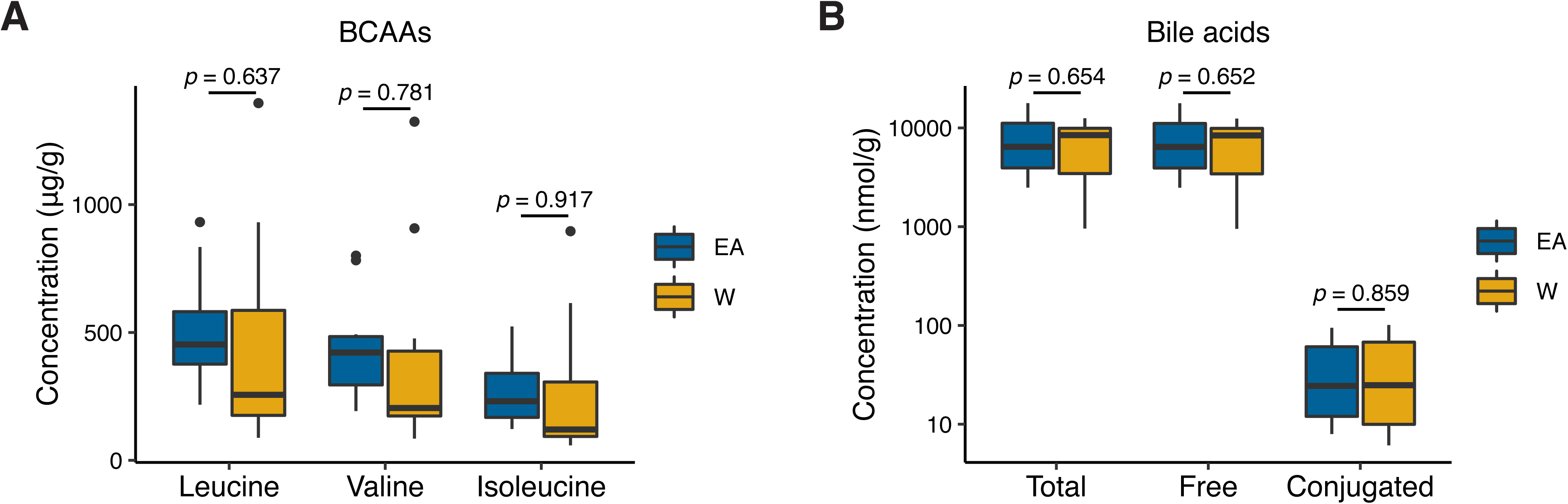
Stool concentrations of branched-chain amino acids and bile acids are comparable between East Asian and White subjects. We did not detect a significant difference in the concentrations of BCAAs **(A)** or bile acids **(B)** between EA (n=10) and W (n=10) stool samples. Statistical analyses performed using Wilcoxon rank-sum tests.

**Figure S3.**
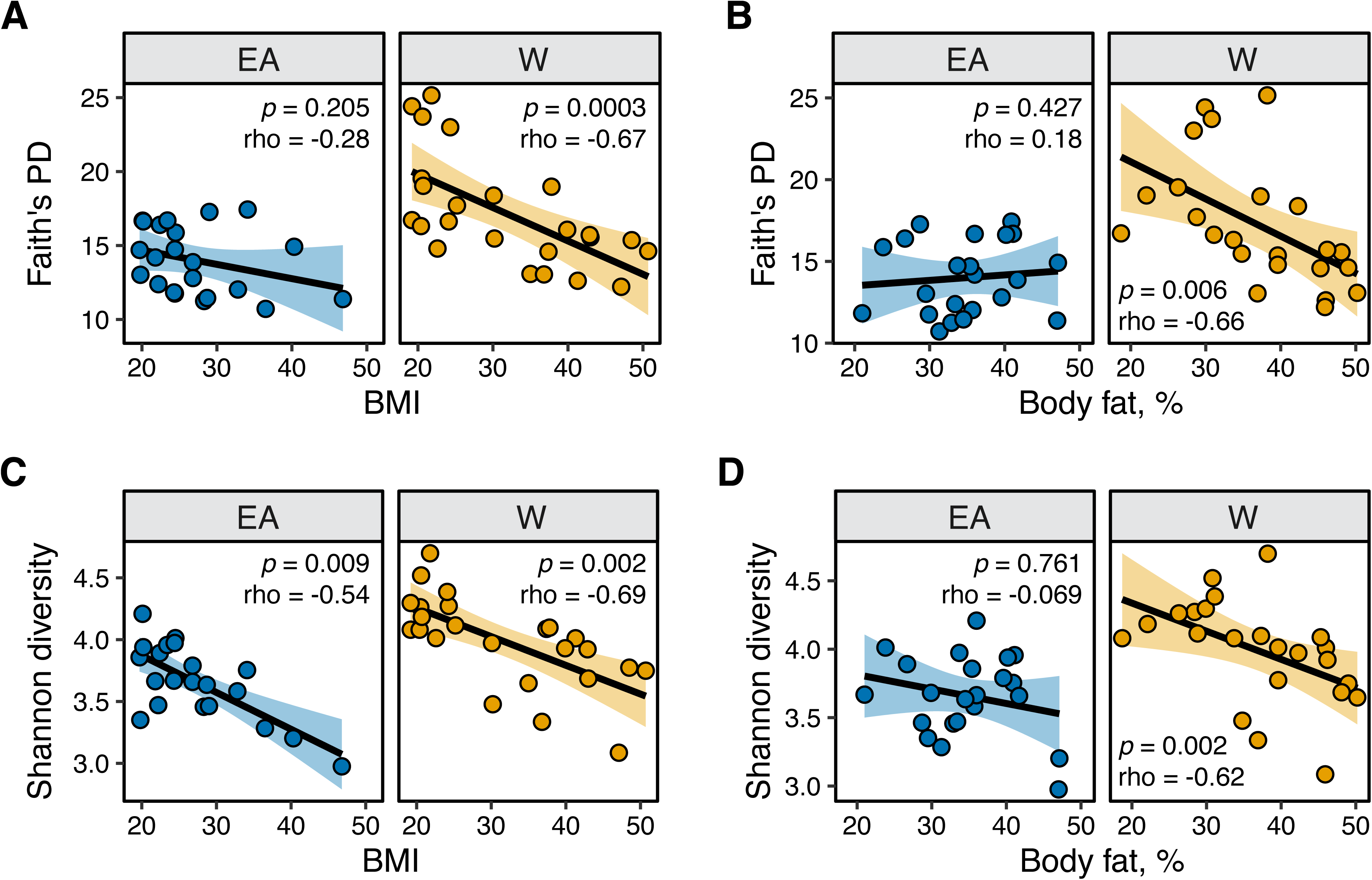
Microbial diversity metrics are consistently and negatively correlated with metabolic parameters in White individuals. **(A,B)** Faith’s diversity is significantly correlated with (A) BMI and (B) percent body fat in W but not EA individuals. **(C,D)** Shannon diversity is significantly correlated with (C) BMI in both W and EA individuals, and with (D) percent body fat in W but not EA individuals. Spearman rank correlation coefficients and *p*-values are shown for each graph (n=12 EA lean, 10 EA obese, 11 W lean, and 13 W obese individuals).

**Figure S4.**
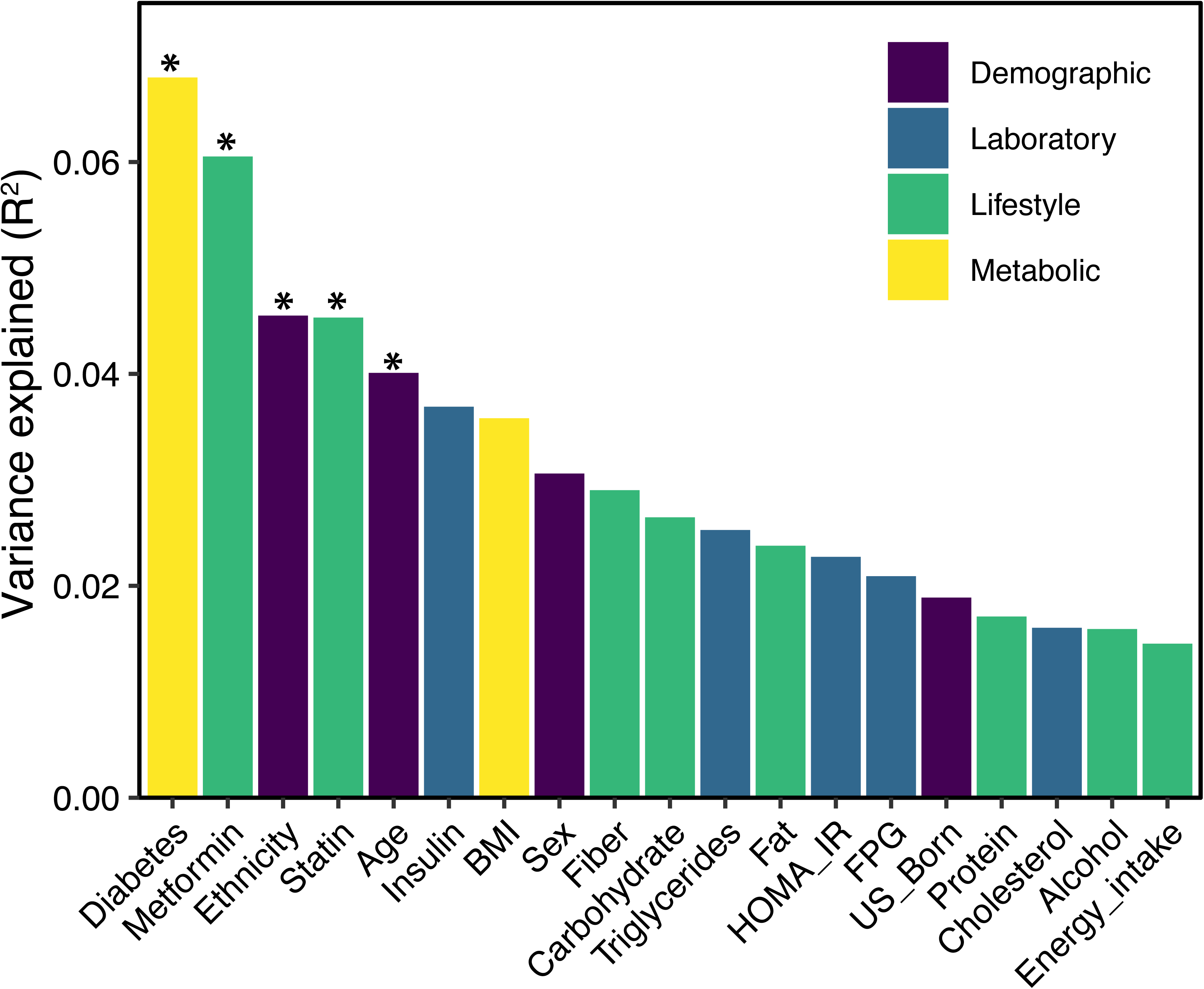
Variables associated with variance in microbial 16S-Seq data. PERMANOVA calculations for metadata variables on the x-axis with relation to variance in PhILR-transformed 16S-Seq data were calculated using the vegan package “adonis”. The resulting effect size is plotted on the y-axis. Variables are colored by type (legend). Asterisks indicate the variables that were statistically significant (*p*<0.05, ADONIS).

**Figure S5.**
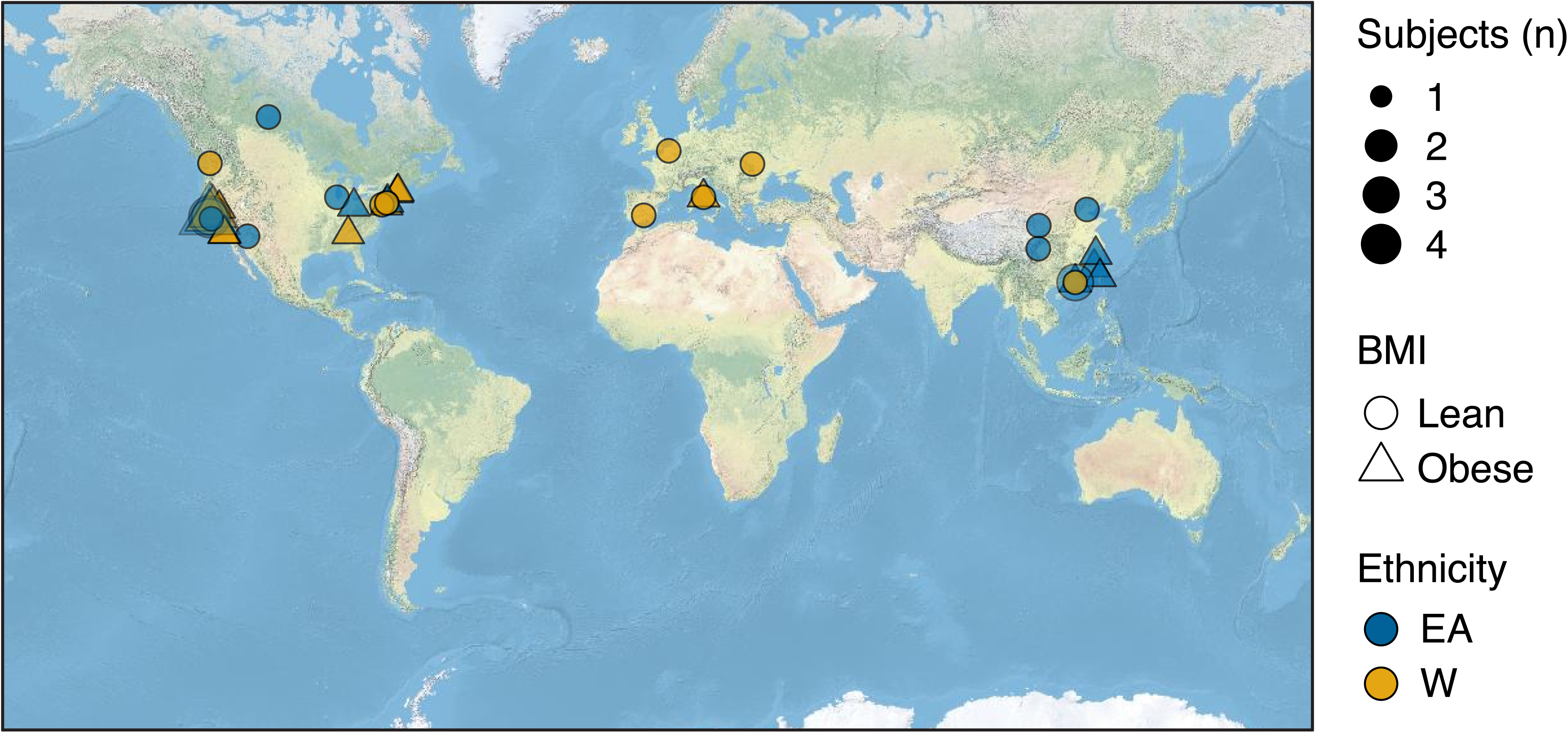
Birth location of subjects. Symbols representing subjects’ birth locations are plotted on a world map. The size, shape and color of the symbols represent the number, BMI and ethnicity of subjects at each location.

**Figure S6.**
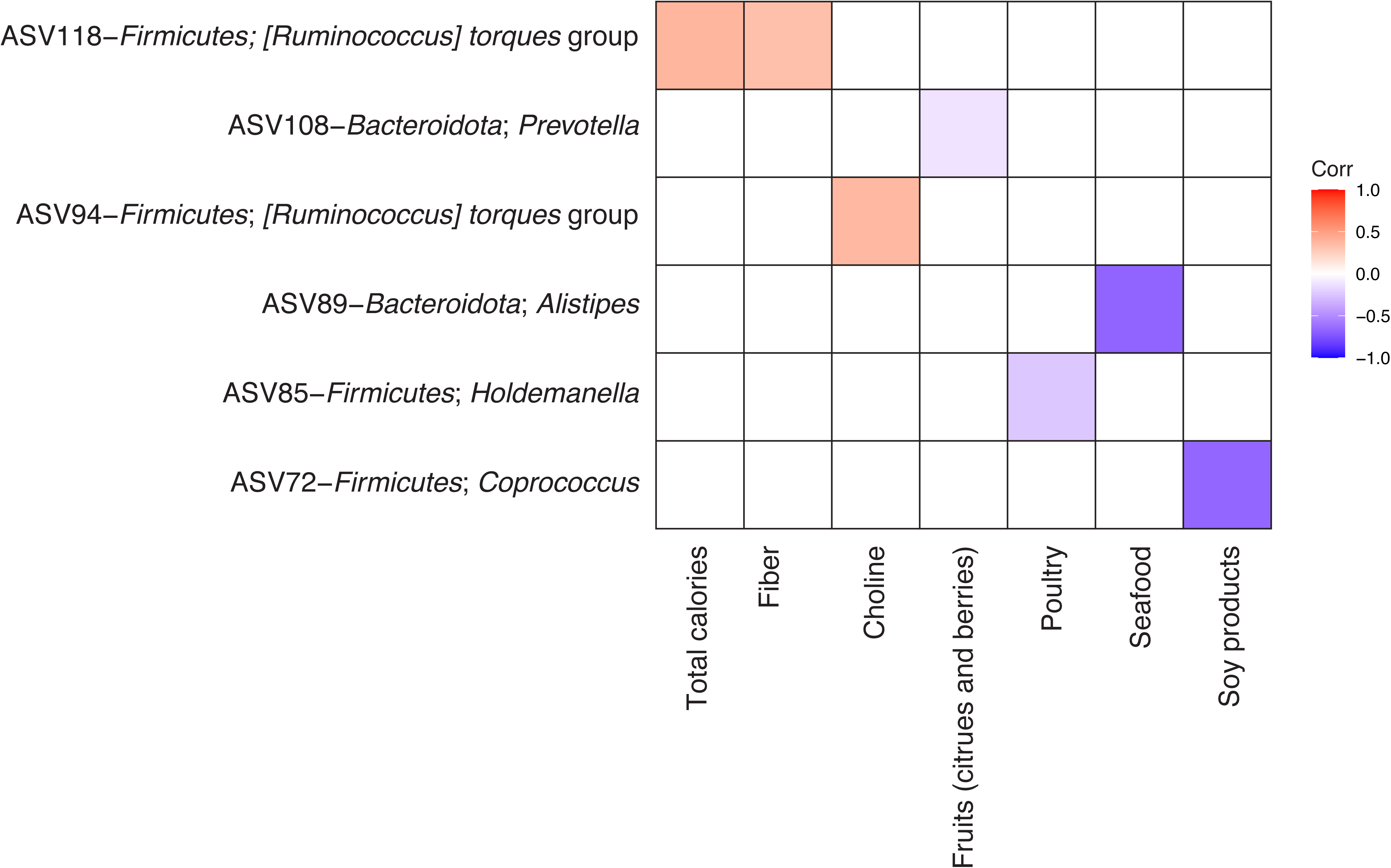
Identification of ASVs associated with short-term dietary intake. Spearman’s correlation was calculated between all 16S-seq ASVs and ASA24 data for lean W subjects. Colored boxes indicate correlations that met our FDR<0.1 cutoff and the direction and intensity of the Spearman’s correlation are shown with correlation color indicated in the figure legend. In lean EA subjects there was a negative correlation between “4:0 Butanoic acid” (g) and ASV46−Firmicutes−Megamonas that was not found in lean W subjects.

**Figure S7.**
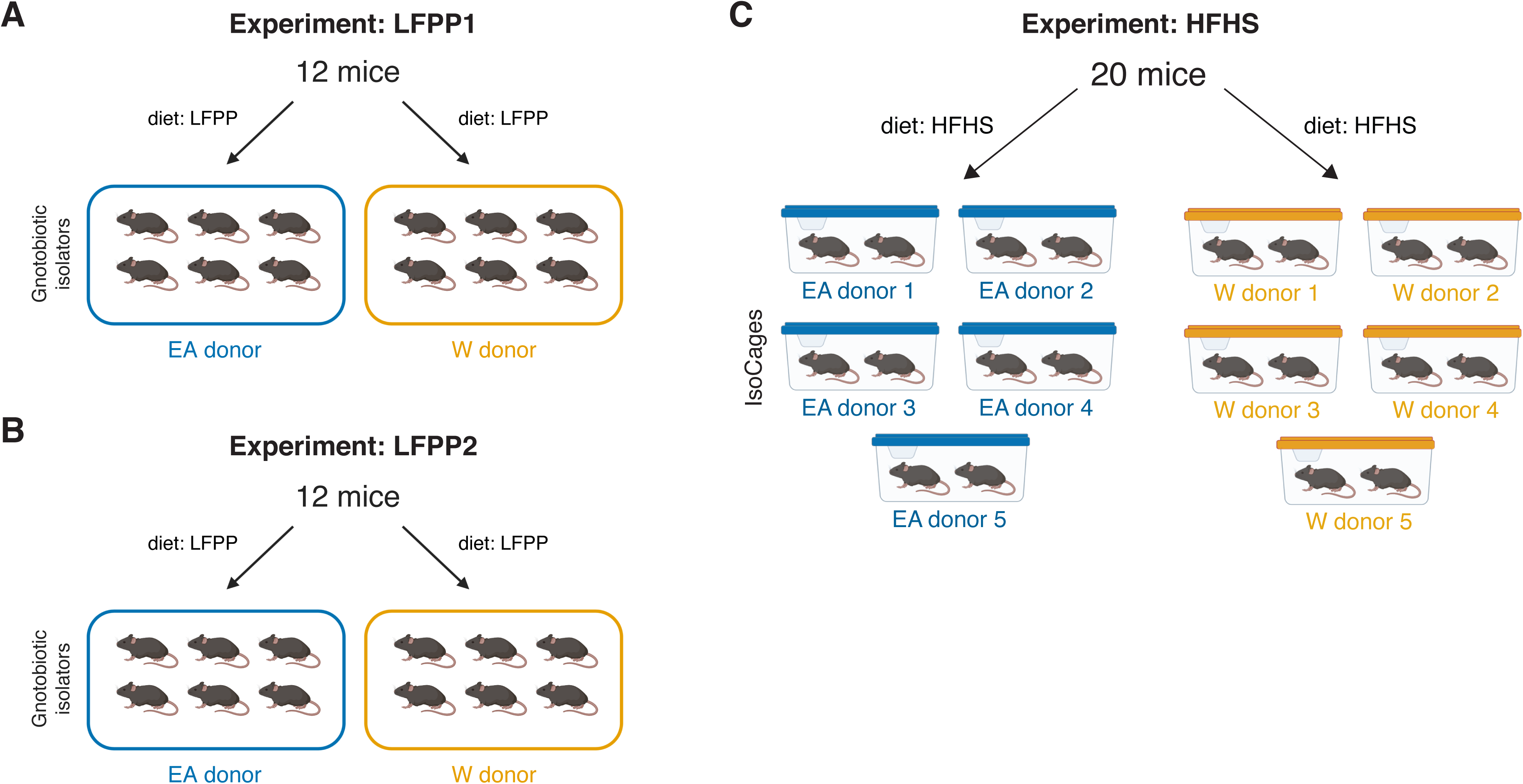
Experimental designs for gnotobiotic experiments. **(A)** LFPP1 experiment: Germ-free mice fed a LFPP diet received an aliquot of stool from a donor of either ethnicity and monitored over 6 weeks (per donor n=6 recipient mice, 1 isolator, 2 cages). **(B)** LFPP2 experiment: Same experimental design as LFPP1 but colonization time was shortened to 3 weeks and two new donor samples were used (per donor n=6 recipient mice, 1 isolator, 2 cages). (**C**) HFHS experiment: 5 lean EA and 5 W donors stool microbial communities were transplanted into 20 germ-free recipient mice fed a HFHS diet for 4 weeks prior to colonization and maintained on diet for another 3 weeks post-transplantation (per donor n=2 recipient mice, 1 IsoCage).

**Figure S8.**
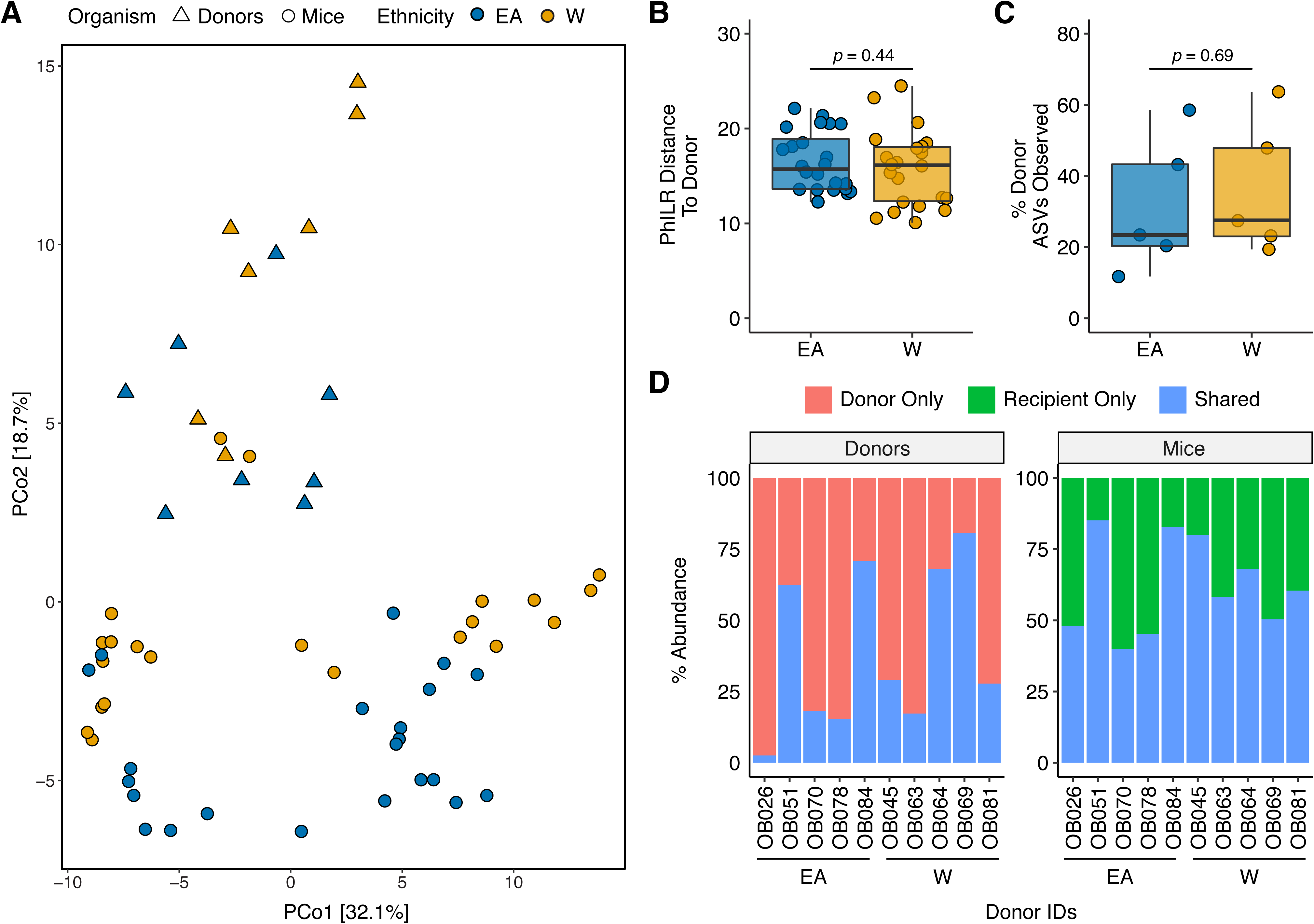
Engraftment efficiency is comparable between donor groups. **(A)** PhILR transformed 16S-Seq data is plotted for gnotobiotic donor slurries (triangles) and recipient mice (circles) and colored by ethnicity (EA, blue; W, orange). See donor metadata in **Tables S2,S4**. There was no significant difference in **(B)** the PhILR distance of recipient mice to their respective donors or **(C)** the proportion of donor ASVs detected within recipient mice between groups (Wilcoxon rank-sum test). **(D)** Relative abundance of shared and unique ASVs in each donor and the corresponding recipient mice.

**Figure S9.**
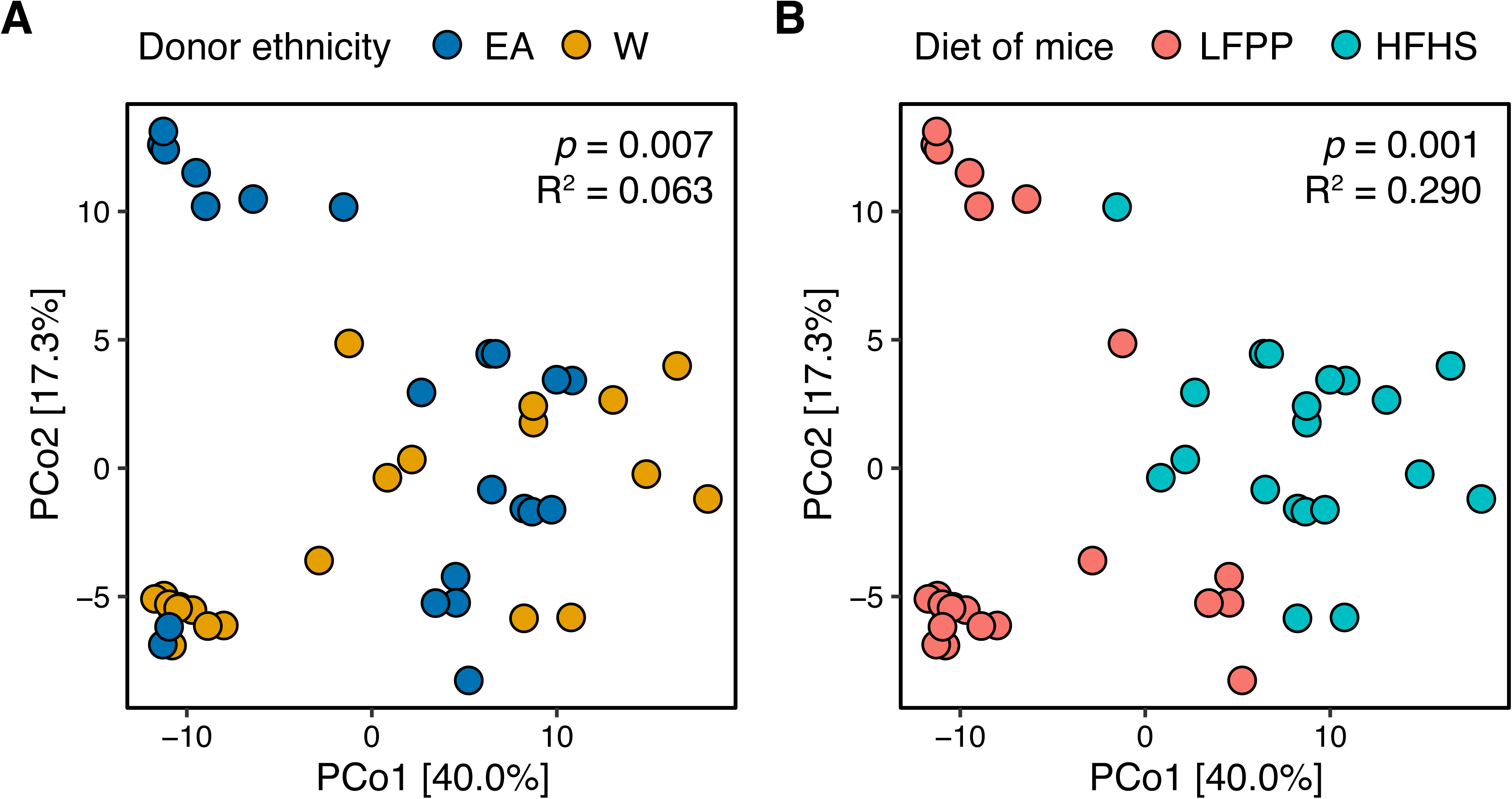
Combined analysis of recipient mice reveals significant associations with donor ethnicity and recipient diet. A PhILR PCoA is plotted based on 16S-Seq data from all gnotobiotic experiments. Individual mice are colored by **(A)** donor ethnicity or **(B)** the recipient’s diet. Both ethnicity and diet were statistically significant contributors to variance (ADONIS *p*-values and estimated variance displayed). We also observed an interaction between diet and ethnicity in this model (*p*=0.016, R^2^=0.047, ADONIS).

## LIST OF ABBREVIATIONS

16S-seq: 16S rRNA gene sequencing
ASA24: Automated Self Administered 24-Hour Dietary Assessment Tool
ASV: Amplicon Sequence Variant
BCAA: branched chain amino acid
BMI: body mass index
DHQIII: Diet History Questionnaire III
EA: East Asian
HFHS: high-fat, high-sugar
LFPP: low-fat, plant-polysaccharide rich
SCFA: short-chain fatty acid
W: White
ZCTA: ZIP Code Tabulation Area

## DECLARATIONS

### Ethics approval and consent to participate

Human stool samples were collected as part of a multi-ethnic clinical cohort study termed Inflammation, Diabetes, Ethnicity and Obesity (ClinicalTrials.gov identifier NCT03022682), consisting of 25- to 65-year-old men and women residing in Northern California and recruited from medical and surgical clinics at UCSF and the Zuckerberg San Francisco General Hospital, or through local public advertisements. The host phenotypic data from this cohort have been described in detail [18, 25]. Informed consent was provided for all subjects participating in the study, which was approved by the UCSF Institutional Review Board. Protocols for all experiments involving mice were approved by the University of California, San Francisco Institutional Animal Care and Use Committee, and performed accordingly.

### Consent for publication

Not applicable.

### Availability of data and materials

All 16S-seq and metagenomic sequencing data generated in the preparation of this manuscript have been deposited in NCBI’s Sequence Read Archive under accession number PRJNA665061. Metabolomics results and metadata are available within this manuscript (**Tables S2, S3, S7,** and **S8**). Code for our manuscript will be uploaded to GitHub (https://github.com/turnbaughlab/2021_IDEO).

### Authors’ contributions

PJT and SKK designed the study. DLA and SKK recruited the IDEO cohort. QYA and DLA performed all gnotobiotic mouse experiments. QYA and EB performed DNA extractions and library preparations for sequencing. QYA and VU conducted all bioinformatic analyses including analysis of microbiome, geographic, and dietary datasets. JEB wrote the bioinformatic pipelines for processing 16S-seq and metagenomic data and assisted with data analysis and interpretation. HLL, GW, and EB helped with the annotation and analysis of subjects’ metadata and dietary questionnaires. CN contributed code to generate the phylogenetic tree used in the supplement. JC and ADP conducted and analyzed metabolomics data. PJT, QYA, and VU wrote the manuscript and prepared the figures with input from all co-authors.

## Acknowledgements

We thank Jessie Turnbaugh and the other UCSF Gnotobiotics Core Facility staff and members of the Koliwad lab for help with the gnotobiotic mouse experiments. We thank Dr. Philip B. Smith from the Penn State Metabolomics Facility. We also thank the CZ Biohub Sequencing Platform for sequencing support, as well as all the subjects who participated in this study.

## REFERENCES

1. Yatsunenko T, Rey FE, Manary MJ, Trehan I, Dominguez-Bello MG, Contreras M, et al. Human gut microbiome viewed across age and geography. Nature. Nature Publishing Group; 2012;486:222–7.

2. Hehemann J-H, Correc G, Barbeyron T, Helbert W, Czjzek M, Michel G. Transfer of carbohydrate-active enzymes from marine bacteria to Japanese gut microbiota. Nature. Nature Publishing Group; 2010;464:908–12.

3. Vangay P, Johnson AJ, Ward TL, Al-Ghalith GA, Shields-Cutler RR, Hillmann BM, et al. US Immigration Westernizes the Human Gut Microbiome. Cell. Nature Publishing Group; 2018;175:962–72.e10.

4. De Filippo C, Cavalieri D, Di Paola M, Ramazzotti M, Poullet JB, Massart S, et al. Impact of diet in shaping gut microbiota revealed by a comparative study in children from Europe and rural Africa. Proc Natl Acad Sci USA. 2010;107:14691–6.

5. Devoto AE, Santini JM, Olm MR, Anantharaman K, Munk P, Tung J, et al. Megaphages infect Prevotella and variants are widespread in gut microbiomes. Nat Microbiol. 2019;4:693– 700.

6. David LA, Maurice CF, Carmody RN, Gootenberg DB, Button JE, Wolfe BE, et al. Diet rapidly and reproducibly alters the human gut microbiome. Nature. 2014;505:559–63.

7. Carmody RN, Gerber GK, Luevano JM Jr, Gatti DM, Somes L, Svenson KL, et al. Diet dominates host genotype in shaping the murine gut microbiota. Cell Host Microbe. 2015;17:72– 84.

8. Gehrig JL, Venkatesh S, Chang H-W, Hibberd MC, Kung VL, Cheng J, et al. Effects of microbiota-directed foods in gnotobiotic animals and undernourished children. Science. 2019;365.

9. Bisanz JE, Upadhyay V, Turnbaugh JA, Ly K, Turnbaugh PJ. Meta-Analysis Reveals Reproducible Gut Microbiome Alterations in Response to a High-Fat Diet. Cell Host Microbe. 2019;26:265–72.e4.

10. Khine WWT, Zhang Y, Goie GJY, Wong MS, Liong M, Lee YY, et al. Gut microbiome of pre-adolescent children of two ethnicities residing in three distant cities. Sci Rep. 2019;9:7831.

11. Deschasaux M, Bouter KE, Prodan A, Levin E, Groen AK, Herrema H, et al. Depicting the composition of gut microbiota in a population with varied ethnic origins but shared geography. Nat Med. 2018;24:1526–31.

12. Xu J, Lawley B, Wong G, Otal A, Chen L, Ying TJ, et al. Ethnic diversity in infant gut microbiota is apparent before the introduction of complementary diets. Gut Microbes. 2020;11:1362–73.

13. Brooks AW, Priya S, Blekhman R, Bordenstein SR. Gut microbiota diversity across ethnicities in the United States. PLoS Biol. 2018;16:e2006842.

14. Sordillo JE, Zhou Y, McGeachie MJ, Ziniti J, Lange N, Laranjo N, et al. Factors influencing the infant gut microbiome at age 3-6 months: Findings from the ethnically diverse Vitamin D Antenatal Asthma Reduction Trial (VDAART). J Allergy Clin Immunol. 2017;139:482–91.e14.

15. Zheng W, McLerran DF, Rolland B, Zhang X, Inoue M, Matsuo K, et al. Association between body-mass index and risk of death in more than 1 million Asians. N Engl J Med. 2011;364:719–29.

16. Gu D, He J, Duan X, Reynolds K, Wu X, Chen J, et al. Body weight and mortality among men and women in China. JAMA. 2006;295:776–83.

17. Jih J, Mukherjea A, Vittinghoff E, Nguyen TT, Tsoh JY, Fukuoka Y, et al. Using appropriate body mass index cut points for overweight and obesity among Asian Americans. Prev Med. 2014;65:1–6.

18. Alba DL, Farooq JA, Lin MYC, Schafer AL, Shepherd J, Koliwad SK. Subcutaneous Fat Fibrosis Links Obesity to Insulin Resistance in Chinese Americans. J Clin Endocrinol Metab. 2018;103:3194–204.

19. Xiang K, Wang Y, Zheng T, Jia W, Li J, Chen L, et al. Genome-wide search for type 2 diabetes/impaired glucose homeostasis susceptibility genes in the Chinese: significant linkage to chromosome 6q21-q23 and chromosome 1q21-q24. Diabetes. 2004;53:228–34.

20. Wen J, Rönn T, Olsson A, Yang Z, Lu B, Du Y, et al. Investigation of type 2 diabetes risk alleles support CDKN2A/B, CDKAL1, and TCF7L2 as susceptibility genes in a Han Chinese cohort. PLoS One. journals.plos.org; 2010;5:e9153.

21. Gravel S, Henn BM, Gutenkunst RN, Indap AR, Marth GT, Clark AG, et al. Demographic history and rare allele sharing among human populations. Proceedings of the National Academy of Sciences. 2011;108:11983–8.

22. Ley RE, Turnbaugh PJ, Klein S, Gordon JI. Human gut microbes associated with obesity. Nature. 2006;444:1022–3.

23. Turnbaugh PJ, Hamady M, Yatsunenko T, Cantarel BL, Duncan A, Ley RE, et al. A core gut microbiome in obese and lean twins. Nature. 2009;457:480–4.

24. Wu H, Tremaroli V, Schmidt C, Lundqvist A, Olsson LM, Krämer M, et al. The Gut Microbiota in Prediabetes and Diabetes: A Population-Based Cross-Sectional Study. Cell Metab. 2020;32:379–90.

25. Oguri Y, Shinoda K, Kim H, Alba DL, Bolus WR, Wang Q, et al. CD81 Controls Beige Fat Progenitor Cell Growth and Energy Balance via FAK Signaling. Cell. 2020;182:563–77.e20.

26. Hsu WC, Araneta MRG, Kanaya AM, Chiang JL, Fujimoto W. BMI cut points to identify at-risk Asian Americans for type 2 diabetes screening. Diabetes Care. 2015;38:150–8.

27. WHO Expert Consultation. Appropriate body-mass index for Asian populations and its implications for policy and intervention strategies. Lancet. 2004;363:157–63.

28. McClung HL, Ptomey LT, Shook RP, Aggarwal A, Gorczyca AM, Sazonov ES, et al. Dietary Intake and Physical Activity Assessment: Current Tools, Techniques, and Technologies for Use in Adult Populations. Am J Prev Med. 2018;55:e93–104.

29. American Diabetes Association. 2. Classification and Diagnosis of Diabetes: Standards of Medical Care in Diabetes—2019. Diabetes Care. 2019;42:S13–28.

30. Friedewald WT, Levy RI, Fredrickson DS. Estimation of the concentration of low-density lipoprotein cholesterol in plasma, without use of the preparative ultracentrifuge. Clin Chem. 1972;18:499–502.

31. Matthews DR, Hosker JP, Rudenski AS, Naylor BA, Treacher DF, Turner RC. Homeostasis model assessment: insulin resistance and beta-cell function from fasting plasma glucose and insulin concentrations in man. Diabetologia. 1985;28:412–9.

32. Kaul S, Rothney MP, Peters DM, Wacker WK, Davis CE, Shapiro MD, et al. Dual-Energy X-Ray Absorptiometry for Quantification of Visceral Fat. Obesity. 2012. p. 1313–8.

33. Bredella MA, Gill CM, Keating LK, Torriani M, Anderson EJ, Punyanitya M, et al. Assessment of abdominal fat compartments using DXA in premenopausal women from anorexia nervosa to morbid obesity. Obesity. 2013;21:2458–64.

34. Neeland IJ, Grundy SM, Li X, Adams-Huet B, Vega GL. Comparison of visceral fat mass measurement by dual-X-ray absorptiometry and magnetic resonance imaging in a multiethnic cohort: the Dallas Heart Study. Nutr Diabetes. 2016;6:e221.

35. Craig CL, Marshall AL, Sjöström M, Bauman AE, Booth ML, Ainsworth BE, et al. International physical activity questionnaire: 12-country reliability and validity. Med Sci Sports Exerc. 2003;35:1381–95.

36. Timon CM, van den Barg R, Blain RJ, Kehoe L, Evans K, Walton J, et al. A review of the design and validation of web- and computer-based 24-h dietary recall tools. Nutrition Research Reviews. 2016. p. 268–80.

37. Park Y, Dodd KW, Kipnis V, Thompson FE, Potischman N, Schoeller DA, et al. Comparison of self-reported dietary intakes from the Automated Self-Administered 24-h recall, 4-d food records, and food-frequency questionnaires against recovery biomarkers. The American Journal of Clinical Nutrition. 2018. p. 80–93.

38. Millen AE, Midthune D, Thompson FE, Kipnis V, Subar AF. The National Cancer Institute diet history questionnaire: validation of pyramid food servings. Am J Epidemiol. 2006;163:279– 88.

39. Diet History Questionnaire III (DHQ III) . [cited 2020 Sep 15]. Available from: https://epi.grants.cancer.gov/dhq3/

40. Caporaso JG, Lauber CL, Walters WA, Berg-Lyons D, Huntley J, Fierer N, et al. Ultra-high-throughput microbial community analysis on the Illumina HiSeq and MiSeq platforms. ISME J. 2012;6:1621–4.

41. Gohl DM, Vangay P, Garbe J, MacLean A, Hauge A, Becker A, et al. Systematic improvement of amplicon marker gene methods for increased accuracy in microbiome studies. Nat Biotechnol. 2016;34:942–9.

42. Callahan BJ, McMurdie PJ, Rosen MJ, Han AW, Johnson AJA, Holmes SP. DADA2: High-resolution sample inference from Illumina amplicon data. Nat Methods. 2016;13:581–3.

43. Wang Q, Garrity GM, Tiedje JM, Cole JR. Naive Bayesian classifier for rapid assignment of rRNA sequences into the new bacterial taxonomy. Appl Environ Microbiol. 2007;73:5261–7.

44. Dixon P. VEGAN, a package of R functions for community ecology. J Veg Sci. 2003;14:927–30.

45. Kembel SW, Cowan PD, Helmus MR, Cornwell WK, Morlon H, Ackerly DD, et al. Picante: R tools for integrating phylogenies and ecology. Bioinformatics. 2010;26:1463–4.

46. Silverman JD, Washburne AD, Mukherjee S, David LA. A phylogenetic transform enhances analysis of compositional microbiota data. Elife. 2017;6.

47. Paradis E, Claude J, Strimmer K. APE: analyses of phylogenetics and evolution in R language. Bioinformatics. 2004;20:289–90.

48. Bisanz J. MicrobeR v 0.3.2: Handy functions for microbiome analysis in R. GitHub. 2017; Available from: github.com/jbisanz/MicrobeR.

49. Martín-Fernández J-A, Hron K, Templ M, Filzmoser P, Palarea-Albaladejo J. Bayesian-multiplicative treatment of count zeros in compositional data sets. Stat Modelling. 2015;15:134– 58.

50. Fernandes AD, Macklaim JM, Linn TG, Reid G, Gloor GB. ANOVA-like differential gene expression analysis of single-organism and meta-RNA-seq. PLoS One. 2013;8:e67019.

51. Benjamini Y, Hochberg Y. Controlling the false discovery rate: a practical and powerful approach to multiple testing. J R Stat Soc Series B Stat Methodol. Wiley; 1995;57:289–300.

52. Kassambara A, Kassambara MA. Package “ggcorrplot.” R package version 0.1.1. mran.microsoft.com; 2019;3. Available from: https://mran.microsoft.com/snapshot/2018-06-22/web/packages/ggcorrplot/ggcorrplot.pdf

53. Liaw A, Wiener M, Others. Classification and regression by randomForest. R news. 2002;2:18–22.

54. Robin X, Turck N, Hainard A, Tiberti N, Lisacek F, Sanchez J-C, et al. pROC: an open- source package for R and S+ to analyze and compare ROC curves. BMC Bioinformatics. 2011. p. 77.

55. Sing T, Sander O, Beerenwinkel N, Lengauer T. ROCR: visualizing classifier performance in R . Bioinformatics. 2005. p. 7881.

56. Chen S, Zhou Y, Chen Y, Gu J. fastp: an ultra-fast all-in-one FASTQ preprocessor. Bioinformatics. 2018;34:i884–90.

57. Uritskiy GV, DiRuggiero J, Taylor J. MetaWRAP-a flexible pipeline for genome-resolved metagenomic data analysis. Microbiome. 2018;6:158.

58. Segata N, Waldron L, Ballarini A, Narasimhan V, Jousson O, Huttenhower C. Metagenomic microbial community profiling using unique clade-specific marker genes. Nat Methods. 2012;9:811–4.

59. Franzosa EA, McIver LJ, Rahnavard G, Thompson LR, Schirmer M, Weingart G, et al. Species-level functional profiling of metagenomes and metatranscriptomes. Nat Methods. 2018;15:962–8.

60. Nayfach S, Pollard KS. Average genome size estimation improves comparative metagenomics and sheds light on the functional ecology of the human microbiome. Genome Biol. 2015;16:51.

61. Cai J, Zhang J, Tian Y, Zhang L, Hatzakis E, Krausz KW, et al. Orthogonal comparison of GC-MS and 1H NMR spectroscopy for short chain fatty acid quantitation. Anal Chem. 2017;89:7900–6.

62. Zheng X, Qiu Y, Zhong W, Baxter S, Su M, Li Q, et al. A targeted metabolomic protocol for short-chain fatty acids and branched-chain amino acids. Metabolomics. 2013;9:818–27.

63. Sarafian MH, Lewis MR, Pechlivanis A, Ralphs S, McPhail MJW, Patel VC, et al. Bile acid profiling and quantification in biofluids using ultra-performance liquid chromatography tandem mass spectrometry. Anal Chem. 2015;87:9662–70.

64. Kahle D, Wickham H. ggmap: Spatial Visualization with ggplot2. The R Journal. 2013;5:144–61.

65. Fellows I, Stotz JP. OpenStreetMap: Access to open street map raster images. R Package Version, 0.3.3. 2016.

66. Wallace JR. Imap: Interactive mapping. R package version 1.32. R Found Stat Comput; 2012.

67. Kassambara A. ggpubr:“ggplot2” based publication ready plots. R package version 0.1. 2018;7.

68. Cheng J, Karambelkar B, Xie Y. Leaflet: Create interactive web maps with the Javascript leaflet library. 2018;2.

69. Maechler M, Rousseeuw P, Struyf A, Hubert M, Hornik K, Others. Cluster: cluster analysis basics and extensions. 2012;1:56.

70. Wickham H, Bryan J. readxl: Read Excel Files. R package version 1.0.0. URL: https://CRAN R-project.org/package=readxl. 2017.

71. Krijthe JH. Rtsne: T-distributed stochastic neighbor embedding using Barnes-Hut implementation. R package version 0.13, URL: https://github com/jkrijthe/Rtsne. 2015.

72. Oksanen J, Blanchet FG, Kindt R, Legendre P, Minchin PR, O’hara RB, et al. Community ecology package. 2013;2–0.

73. Paradis E, Schliep K. ape 5.0: an environment for modern phylogenetics and evolutionary analyses in R. Bioinformatics. 2019.

74. Walker K. Tigris: Load census TIGER/Line Shapefiles. R package version 0.7. 2018.

75. Kuznetsova A, Brockhoff PB, Christensen RHB, Others. lmerTest package: tests in linear mixed effects models. J Stat Softw. 2017;82:1–26.

76. Bisanz JE. qiime2R: Importing QIIME2 artifacts and associated data into R sessions. Version 0.99. 2018;13.

77. Yutani H. gghighlight: Highlight Lines and Points in “ggplot2.” Manual available online at http://CRAN.R-project org/package=gghighlight. 2018.

78. McMurdie PJ, Holmes S. phyloseq: an R package for reproducible interactive analysis and graphics of microbiome census data. PLoS One. 2013;8:e61217.

79. Firke S. Janitor: Simple tools for examining and cleaning dirty data. 2018;1.

80. Rich B. table1: Tables of Descriptive Statistics in HTML. R package version 1.2. 2020. Available from: https://CRAN.R-project.org/package=table1.

81. Wickham H. ggplot2: Elegant Graphics for Data Analysis. 2016. Available from: https://ggplot2.tidyverse.org.

82. Le Chatelier E, Nielsen T, Qin J, Prifti E, Hildebrand F, Falony G, et al. Richness of human gut microbiome correlates with metabolic markers. Nature. 2013;500:541–6.

83. Falony G, Joossens M, Vieira-Silva S, Wang J, Darzi Y, Faust K, et al. Population-level analysis of gut microbiome variation. Science. 2016;352:560–4.

84. Forslund K, Hildebrand F, Nielsen T, Falony G, Le Chatelier E, Sunagawa S, et al. Disentangling type 2 diabetes and metformin treatment signatures in the human gut microbiota. Nature. 2015;528:262–6.

85. Ghosh TS, Das M, Jeffery IB, O’Toole PW. Adjusting for age improves identification of gut microbiome alterations in multiple diseases. Elife. 2020;9.

86. Wu H, Esteve E, Tremaroli V, Khan MT, Caesar R, Mannerås-Holm L, et al. Metformin alters the gut microbiome of individuals with treatment-naive type 2 diabetes, contributing to the therapeutic effects of the drug. Nat Med. 2017;23:850–8.

87. Vieira-Silva S, Falony G, Belda E, Nielsen T, Aron-Wisnewsky J, Chakaroun R, et al. Statin therapy is associated with lower prevalence of gut microbiota dysbiosis. Nature. 2020;581:310– 5.

88. Turnbaugh PJ, Ridaura VK, Faith JJ, Rey FE, Knight R, Gordon JI. The effect of diet on the human gut microbiome: a metagenomic analysis in humanized gnotobiotic mice. Sci Transl Med. 2009;1:6ra14.

89. Nayak RR, Alexander M, Deshpande I, Stapleton-Gray K, Rimal B, Patterson AD, et al. Methotrexate impacts conserved pathways in diverse human gut bacteria leading to decreased host immune activation. Cell Host Microbe. 2021.

90. Walter J, Armet AM, Finlay BB, Shanahan F. Establishing or Exaggerating Causality for the Gut Microbiome: Lessons from Human Microbiota-Associated Rodents. Cell. 2020;180:221–32.

91. Garduño-Diaz SD, Husain W, Ashkanani F, Khokhar S. Meeting challenges related to the dietary assessment of ethnic minority populations. J Hum Nutr Diet. 2014;27:358–66.

92. Martiny JBH, Bohannan BJM, Brown JH, Colwell RK, Fuhrman JA, Green JL, et al. Microbial biogeography: putting microorganisms on the map. Nat Rev Microbiol. 2006;4:102– 12.

93. Kakar V, Voelz J, Wu J, Franco J. The Visible Host: Does race guide Airbnb rental rates in San Francisco? J Hous Econ. 2018;40:25–40.

94. Basolo A, Hohenadel M, Ang QY, Piaggi P, Heinitz S, Walter M, et al. Effects of underfeeding and oral vancomycin on gut microbiome and nutrient absorption in humans. Nat Med. 2020;26:589–98.

95. Plovier H, Everard A, Druart C, Depommier C, Van Hul M, Geurts L, et al. A purified membrane protein from Akkermansia muciniphila or the pasteurized bacterium improves metabolism in obese and diabetic mice. Nat Med. 2017;23:107–13.

96. Depommier C, Everard A, Druart C, Plovier H, Van Hul M, Vieira-Silva S, et al. Supplementation with Akkermansia muciniphila in overweight and obese human volunteers: a proof-of-concept exploratory study. Nat Med. 2019;25:1096–103.

97. Perry RJ, Peng L, Barry NA, Cline GW, Zhang D, Cardone RL, et al. Acetate mediates a microbiome–brain–β-cell axis to promote metabolic syndrome. Nature. 2016;534:213–7.

98. Turnbaugh PJ, Ley RE, Mahowald MA, Magrini V, Mardis ER, Gordon JI. An obesity-associated gut microbiome with increased capacity for energy harvest. Nature. 2006;444:1027– 31.

99. Johnson EL, Heaver SL, Walters WA, Ley RE. Microbiome and metabolic disease: revisiting the bacterial phylum Bacteroidetes. J Mol Med. 2017;95:1–8.

100. Tirosh A, Calay ES, Tuncman G, Claiborn KC, Inouye KE, Eguchi K, et al. The short-chain fatty acid propionate increases glucagon and FABP4 production, impairing insulin action in mice and humans. Sci Transl Med. 2019;11.

101. Lu Y, Fan C, Li P, Lu Y, Chang X, Qi K. Short Chain Fatty Acids Prevent High-fat-diet- induced Obesity in Mice by Regulating G Protein-coupled Receptors and Gut Microbiota. Sci Rep. 2016;6:37589.

102. Müller M, Hernández MAG, Goossens GH, Reijnders D, Holst JJ, Jocken JWE, et al. Circulating but not faecal short-chain fatty acids are related to insulin sensitivity, lipolysis and GLP-1 concentrations in humans. Sci Rep. 2019;9:12515.

103. Berding K, Donovan SM. Diet Can Impact Microbiota Composition in Children With Autism Spectrum Disorder. Front Neurosci. 2018;12:515.

104. Kim KN, Yao Y, Ju SY. Short Chain Fatty Acids and Fecal Microbiota Abundance in Humans with Obesity: A Systematic Review and Meta-Analysis. Nutrients. 2019;11.

105. Johnson AJ, Vangay P, Al-Ghalith GA, Hillmann BM, Ward TL, Shields-Cutler RR, et al. Daily Sampling Reveals Personalized Diet-Microbiome Associations in Humans. Cell Host Microbe. 2019;25:789–802.e5.

